# 3D ultrastructural study of synapses in the human entorhinal cortex

**DOI:** 10.1101/2020.05.11.088435

**Authors:** M Domínguez-Álvaro, M Montero-Crespo, L Blazquez-Llorca, J DeFelipe, L Alonso-Nanclares

**Affiliations:** Laboratorio Cajal de Circuitos Corticales, Centro de Tecnología Biomédica, Universidad Politécnica de Madrid. Pozuelo de Alarcón, 28223, Madrid, Spain; Instituto Cajal, Consejo Superior de Investigaciones Científicas (CSIC), Avda. Doctor Arce, 37 Madrid, 28002, Spain; Depto. Psicobiología, Facultad de Psicología, Universidad Nacional de Educación a Distancia (UNED), c/Juan del Rosal, 10 Madrid, 28040, Spain; Centro de Investigación Biomédica en Red sobre Enfermedades Neurodegenerativas (CIBERNED), ISCIII, Madrid, Spain

**Keywords:** Cerebral cortex, electron microscopy, FIB-SEM, neuropil, synaptology

## Abstract

The entorhinal cortex (EC) is a brain region that has been shown to be essential for memory functions and spatial navigation. However, detailed 3D synaptic morphology analysis and identification of postsynaptic targets at the ultrastructural level have not been performed before in the human EC. In the present study, we used Focused Ion Beam/Scanning Electron Microscopy (FIB/SEM) to perform a three-dimensional analysis of the synapses in the neuropil of medial EC in layers II and III from human brain autopsies. Specifically, we studied synaptic structural parameters of **3561** synapses, which were fully reconstructed in 3D. We analyzed the synaptic density, 3D spatial distribution, and type (excitatory and inhibitory), as well as the shape and size of each synaptic junction. Moreover, the postsynaptic targets of synapses could be clearly determined. The present work constitutes a detailed description of the synaptic organization of the human EC, which is a necessary step to better understand the functional organization of this region in both health and disease.

**Significance Statement:** The present study represents the first attempt to unveil the detailed synaptic organization of the neuropil of the human entorhinal cortex — a brain region that is essential for memory function and spatial navigation. Using 3D electron microscopy, we have characterized the synaptic morphology and identified the postsynaptic targets of thousands of synapses. The results provide a new, large, quantitative ultrastructure dataset of the synaptic organization of the human entorhinal cortex. These data provide critical information to better understand synaptic functionality in the human brain.

**Highlight:** Estimation of the number of synapses, as well as determination of their type, shapes, sizes and postsynaptic targets, provides critical data to better understand synaptic functionality. This study provides a new, large, quantitative ultrastructure dataset of the synaptic organization of the human entorhinal cortex using 3D electron microscopy.

## Introduction

The entorhinal cortex (EC) is a brain region which is located on the anterior part of the mesial temporal lobe and which has been shown to be essential for memory functions and spatial navigation (reviewed in Schultz et al., 2015). A number of neurodegenerative conditions, including Alzheimer’s disease, have been related to alterations in the EC (Braak and Braak, 1992). In particular, cognitive deficits have been linked to alterations in the upper layers of EC (Van Hoesen et al., 1991; Gomez-Isla et al., 1996).

Data regarding connections from human EC are mostly inferred from rodents and non-human primates. The EC itself is the origin of the perforant pathway (from layers II and III), which provides the largest input source to the hippocampal formation, targeting the ammonic fields (CA) CA1, CA2 and CA3, as well as dentate gyrus (DG) and subiculum. Specifically, the EC layer II neurons project primarily to the DG and CA3, while EC layer III neurons send their axons to the subiculum and CA1 (Insausti and Amaral, 2012; Kondo et al., 2009). These direct projections of the EC towards the subiculum and the hippocampus are essential for the proper functioning of the hippocampal formation. The classic trisynaptic circuit: EC layer II→DG→CA3→CA1 seems to be related to the acquisition of new memories, whereas the pathway between EC and CA1 neurons (monosynaptic) is thought to contribute to the strength of previously established memories (Cohen and Squire, 1980). Once the information has passed through the hippocampus, it returns to the neurons of the deep layers of the EC (V/VI). These neurons project to the upper layers of the EC itself, sending the information back to the cortical association areas. In addition to establishing reciprocal connections with different association cortices, the EC establishes interconnections with subcortical structures such as the amygdala, septal nuclei or the thalamus, as well as with adjacent regions such as perirhinal cortex, parahippocampal cortex or insular cortex (Braak and Braak, 1992; DeFelipe et al., 2007; Duvernoy, 2005; Insausti and Amaral, 2012; Van Hoesen et al., 1991).

In this way, the EC acts as a gateway for sensory information to the hippocampal formation, also filtering the return of sensory information processed by the hippocampus to the different association areas. Thus, in terms of connections, the EC can be considered as an interface between the hippocampal formation and a large variety of association and limbic cortices (Lavenex and Amaral, 2000; Solodkin and Van Hoesen, 1996).

Furthermore, the EC is not a homogeneous region, as it presents a number of subfields along its extent (reviewed in Schultz et al., 2015; Insausti et al., 2017). Whereas the medial EC interconnects with the parahippocampal cortex, the lateral EC interconnects with the perirhinal cortex. Both medial and lateral EC are connected to the hippocampal formation, including DG, CAs and subiculum (reviewed in Schultz et al., 2015). Thus, mapping the EC connectivity may contribute to the understanding of its structural design. One possible approach to decipher EC connectivity is its analysis at the ultrastructural level, using electron microscopy (EM), to map true synaptic contacts (or synapses).

In the present study, we used Focused Ion Beam/Scanning Electron Microscopy (FIB/SEM) to perform a three-dimensional (3D) analysis of the synapses found in the neuropil of medial EC in layers II and III from ‘normal’ human brain autopsies (subjects with no recorded neurological or psychiatric alterations) with a short postmortem delay (of less than 3.5h). FIB/SEM has already yielded excellent results in human brain samples (Domínguez-Álvaro et al., 2018, 2019; Montero-Crespo et al., 2020). Specifically, we studied a variety of synaptic structural parameters of **3**,**561** synapses, which were fully reconstructed in 3D. In particular, we analyzed the synaptic density, 3D spatial distribution, and type (excitatory and inhibitory), as well as the shape and size of each synaptic junction. Moreover, the postsynaptic targets of **2**,**768** synapses were clearly determined. These detailed morphological data provide quantitative information on the synaptology of this particular brain region and its layers, thereby defining synaptic organization. Additionally, cytoarchitectural characteristics of these EC samples were examined with light microscopy techniques.

Thus, data from the present work constitutes a detailed description of the synaptic organization in superficial layers of the human EC, which is a necessary step to better understand its functional organization in both health and disease.

## Materials and Methods

### Tissue preparation

Human brain tissue was obtained from autopsies (with short post-mortem delays of less than 3.5 hours) from 4 male subjects with no recorded neurological or psychiatric alterations (supplied by Instituto de Neuropatología del IDIBELL - Hospital Universitario de Bellvitge, Barcelona, Spain; Unidad Asociada Neuromax, Laboratorio de Neuroanatomía Humana, Facultad de Medicina, Universidad de Castilla-La Mancha, Albacete, Spain; and the Laboratorio Cajal de Circuitos Corticales UPM-CSIC, Madrid, Spain) (Table 1). The sampling procedure was approved by the Institutional Ethical Committee. Tissue from some of these human brains has been used in previous studies (Domínguez-Álvaro et al., 2018, 2019; Montero-Crespo et al., 2020).

**Table 1.** Clinical and neuropsychological information. None of the four subjects had recorded neurological or psychiatric alterations.

| **Case** | **Sex** | **Age**<br>**(years)** | **Cause of death** | **Postmortem**<br>**delay (h)** | **Neuropsychological**<br>**diagnosis** |
| --- | --- | --- | --- | --- | --- |
| **AB1** | Male | 45 | Lung cancer | <1 | No neurological<br>alterations |
| **AB3** | Male | 53 | Bladder carcinoma | 3.5 | No neurological<br>alterations |
| **IF10** | Male | 66 | Bronchopneumonia<br>and cardiac failure | 2 | No neurological<br>alterations |
| **M16** | Male | 40 | Traffic accident | 3 | No neurological<br>alterations |

Upon removal, brain tissue was fixed in cold 4% paraformaldehyde (Sigma-Aldrich, St Louis, MO, USA) in 0.1M sodium phosphate buffer (PB; Panreac, 131965, Spain), pH 7.4, for 24–48h. After fixation, the tissue was washed in PB and sectioned coronally in a vibratome (150μm thickness; Vibratome Sectioning System, VT1200S Vibratome, Leica Biosystems, Germany). Sections containing EC were selected and processed for Nissl-staining and immunocytochemistry to determine cytoarchitecture (Fig. 1).

**Figure 1.**
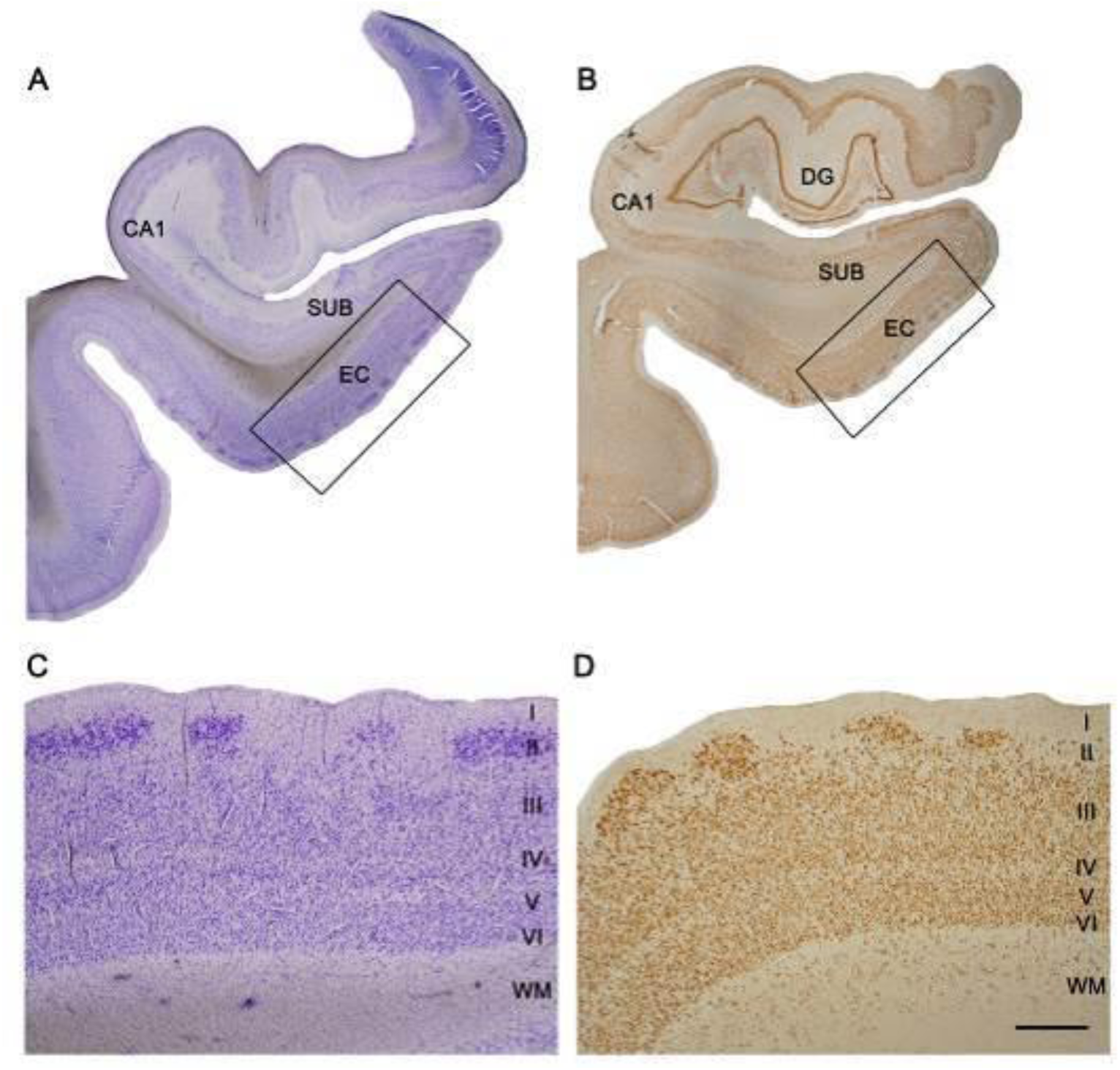
Coronal sections of the human hippocampal formation. (**A, B**) Low-power photographs showing the human EC (boxed areas). **(C, D)** Higher magnification of the boxed areas in A and B, to show the laminar pattern of EC (layers I to VI are indicated). Sections are stained for Nissl (**A, C**) and immunostained for anti-NeuN (**B, D**). Scale bar (in D): 3mm in panels A and B; 600µm in panels C and D. DG: dentate gyrus; EC: entorhinal cortex; SUB: subiculum; WM: white matter.

### Immunohistochemistry

Selected sections were first rinsed in PB 0.1M, pretreated in 2% H_2_O_2_ for 30 minutes to remove endogenous peroxidase activity, and then incubated for 1h at room temperature in a solution of 3% normal horse serum (for polyclonal antisera and monoclonal antibodies, respectively; Vector Laboratories Inc., Burlingame, CA) and 0.25% Triton-X (Merck, Darmstadt, Germany). Subsequently, sections were incubated for 48h at 4°C in the same solution with mouse anti-NeuN (1:2000; Chemicon; MAB377, Temecula, CA, USA). Sections were then processed with a secondary biotinylated horse anti-mouse IgG antibody (1:200, Vector Laboratories, Burlingame, CA, USA). They were then incubated for 1h in an avidin-biotin peroxidase complex (Vectastain ABC Elite PK6100, Vector) and, finally, with the chromogen 3,3′-diaminobenzidine tetrahydrochloride (DAB; Sigma-Aldrich, St. Louis, MO, USA). Sections were then dehydrated, cleared with xylene and cover-slipped.

### Electron microscopy

EC sections were post-fixed for 24h in a solution containing 2% paraformaldehyde, 2.5% glutaraldehyde (TAAB, G002, UK) and 0.003% CaCl_2_ (Sigma, C-2661-500G, Germany) in sodium cacodylate (Sigma, C0250-500G, Germany) buffer (0.1M). These sections were washed in sodium cacodylate buffer (0.1M) and treated with 1% OsO_4_ (Sigma, O5500, Germany), 0.1% potassium ferrocyanide (Probus, 23345, Spain) and 0.003% CaCl_2_ in sodium cacodylate buffer (0.1M) for 1h at room temperature. After washing in PB, sections were stained with 2% uranyl acetate (EMS, 8473, USA), and then dehydrated and flat-embedded in Araldite (TAAB, E021, UK) for 48h at 60°C (DeFelipe and Fairén, 1993). Embedded sections were glued onto a blank Araldite block and trimmed. Semithin sections (1–2 μm thick) were obtained from the surface of the block and stained with 1% toluidine blue (Merck, 115930, Germany) in 1% sodium borate (Panreac, 141644, Spain). The last semithin section (which corresponds to the section immediately adjacent to the block surface) was examined under light microscope and photographed to accurately locate the neuropil regions to be examined (Fig. 2).

**Figure 2.**
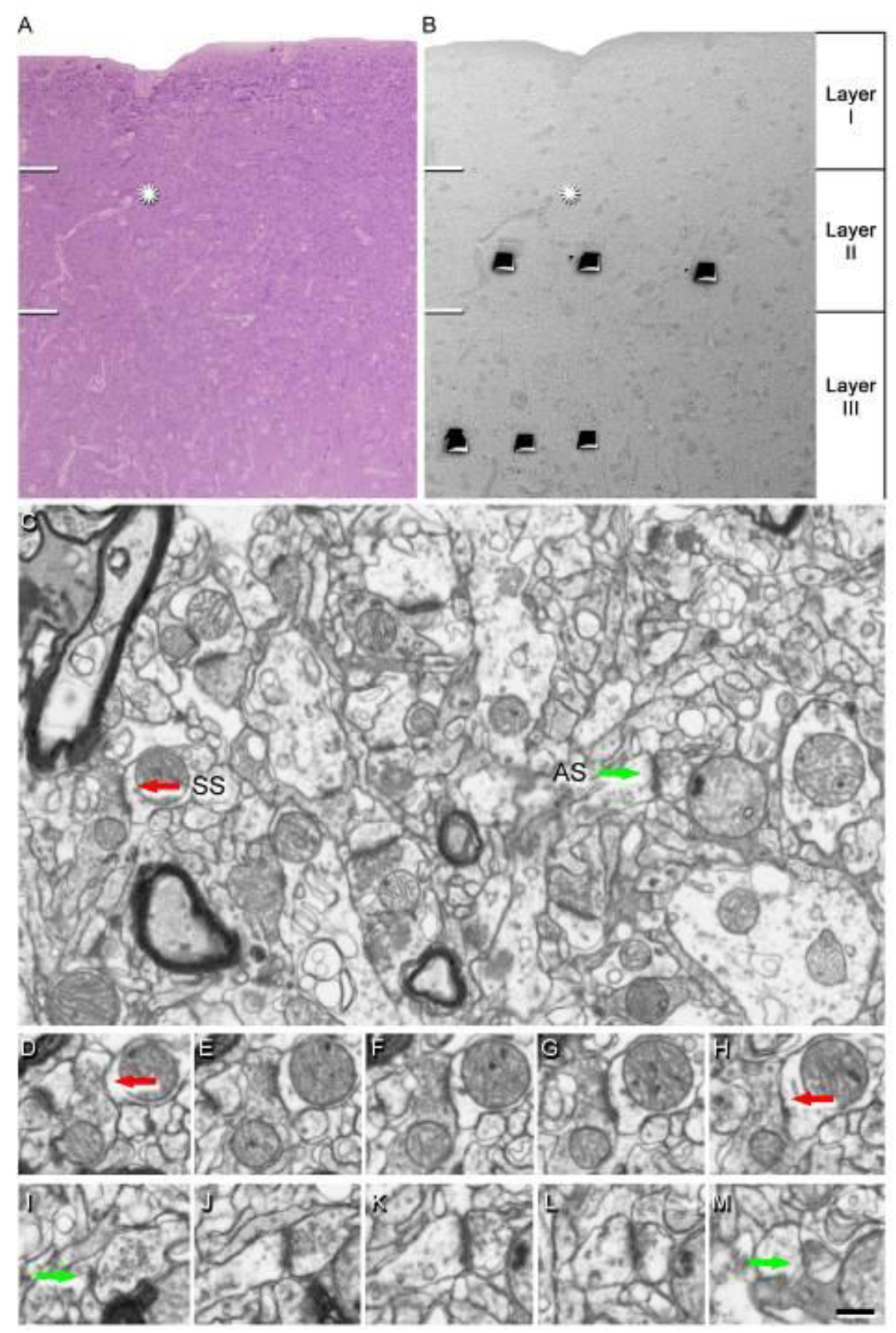
Correlative light/electron microscopy analysis of layer II and III of the EC. Delimitation of layers is based on the staining pattern of 1 µm-thick semithin section, stained with toluidine blue (**A**), which is adjacent to the block for FIB/SEM imaging (**B**). (**B)** SEM image to illustrate the block surface with trenches made in the neuropil (three per layer). Arrows in A and B point to the same blood vessel, showing that the exact location of the region of interest was accurately determined. (**C)** Serial image obtained by FIB/SEM from layer II showing the neuropil, with two synapses indicated (arrows) as examples of asymmetric (AS) and symmetric synapses (SS). Synapse classification was based on the examination of the full sequence of serial images; an SS can be visualized in D−H, and an AS in I−M. Scale bar (in M): 40µm in A; 60µm in B; 1000nm in C; 1300nm in D−M.

### Three-dimensional electron microscopy

The 3D study of the samples was carried out using a dual beam microscope (Crossbeam® 540 electron microscope, Carl Zeiss NTS GmbH, Oberkochen, Germany). This instrument combines a high-resolution field-emission SEM column with a focused gallium ion beam (FIB), which permits removal of thin layers of material from the sample surface on a nanometer scale. As soon as one layer of material (20nm thick) is removed by the FIB, the exposed surface of the sample is imaged by the SEM using the backscattered electron detector. The sequential automated use of FIB milling and SEM imaging allowed us to obtain long series of photographs of a 3D sample of selected regions (Merchán-Pérez et al., 2009). Image resolution in the xy plane was 5nm/pixel. Resolution in the z-axis (section thickness) was 20nm, and image size was 2048 × 1536 pixels. Although the resolution of FIB/SEM images can be increased, we chose these parameters as a compromise solution to obtain a large enough field of view where synaptic junctions could still be clearly identified (Fig. 2) in a reasonable time period that allowed us to have long series of sections (approximately 12 hours per stack of images).

The number of sections per stack from layer II ranged from 244 to 320, which corresponds to a raw volume ranging from 384 to 503 μm^3^ (mean: 454 μm^3^). A total of 12 stacks of images of the neuropil were obtained (three stacks for each of the 4 cases; total volume studied: 5445 μm^3^). For layer III, the number of sections per stack ranged from 268 to 313, corresponding to a corrected volume ranging from 423 to 492 μm^3^ (mean: 456 μm^3^). A total of 12 stacks of neuropil images were obtained (three stacks for each of the 4 cases; total volume studied: 5,466 μm^3^).

A correction in the volume of the stack of images to account for the presence of fixation artifact (i.e., swollen neuronal or glial processes) was applied after quantification with Cavalieri principle (Gundersen et al., 1988). Every FIB/SEM stack was examined and the volume artifact ranged from 3 to 16% of the volume stacks.

All measurements were corrected for tissue shrinkage that occurs during osmication and plastic embedding of the vibratome sections containing the area of interest (Merchán-Pérez et al., 2009). To estimate the shrinkage in our samples, we photographed and measured the vibratome sections with ImageJ (ImageJ 1.51; NIH, USA), both before and after processing for electron microscopy. The values after processing were divided by the values before processing to obtain the volume, area, and linear shrinkage factors (Oorschot et al., 1991), yielding correction factors of 0.90, 0.93, and 0.97, respectively. A total of 24 stacks of images from both layers of the EC were obtained (3 stacks per case and layer for each of the 4 cases, with a total corrected volume studied of 8,592μm^3^; Table 2).

**Table 2.**
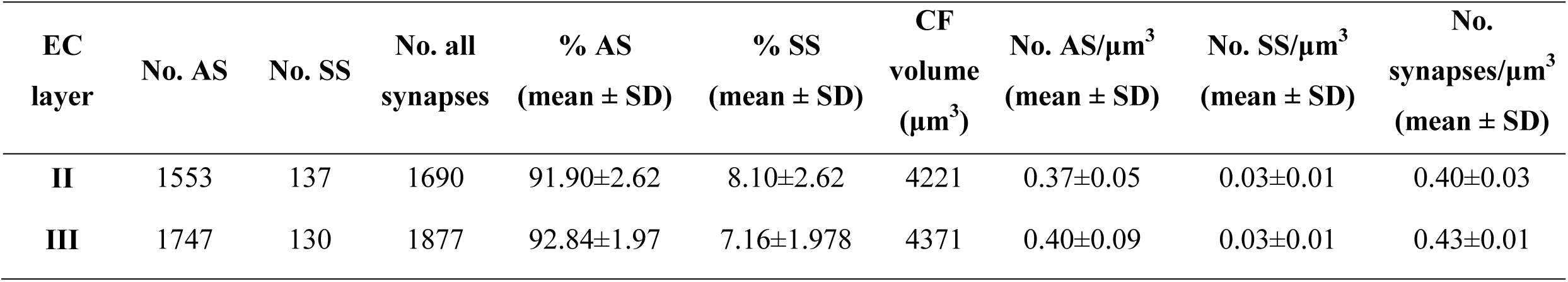
Accumulated data obtained from the ultrastructural analysis of neuropil from layer II and layer III of the EC. All volume data are corrected for fixation artefacts and shrinkage factor. The data for individual cases are shown in Supplementary table 1. AS: asymmetric synapses; CF: counting frame; EC: entorhinal cortex; SD: standard deviation; SS: symmetric synapses.

### Synaptic three-dimensional analysis

Stacks of images obtained by FIB/SEM were analyzed using EspINA software (*EspINA Interactive Neuron Analyzer*, 2.1.9; https://cajalbbp.es/espina/), which allows the segmentation of synapses in the reconstructed 3D volume (for a detailed description of the segmentation algorithm, see Morales et al., 2011; Fig. 3). As previously discussed in (Merchán-Pérez et al., 2009), there is a consensus for classifying cortical synapses into asymmetric synapses (AS; or type I) and symmetric synapses (SS; or type II). The main characteristic distinguishing these synapses is the prominent or thin post-synaptic density, respectively. Nevertheless, in single sections, the synaptic cleft and the pre- and post-synaptic densities are often blurred if the plane of the section does not pass at right angles to the synaptic junction. Since EspINA allows navigation through the stack of images, it was possible to unambiguously identify every synapse as AS or SS, based on the thickness of the PSD (Merchán-Pérez et al., 2009). Synapses with prominent PSDs are classified as AS, while thin PSDs are classified as SS (Gray, 1959; Peters and Palay, 1991; Fig. 2). EspINA provided the number of synapses in a given volume, which allowed the estimation of the number of synapses per volume. EspINA also allowed the application of an unbiased 3D counting frame (CF) to perform direct counting (for details, see Merchán-Pérez et al., 2009).

**Figure 3.**
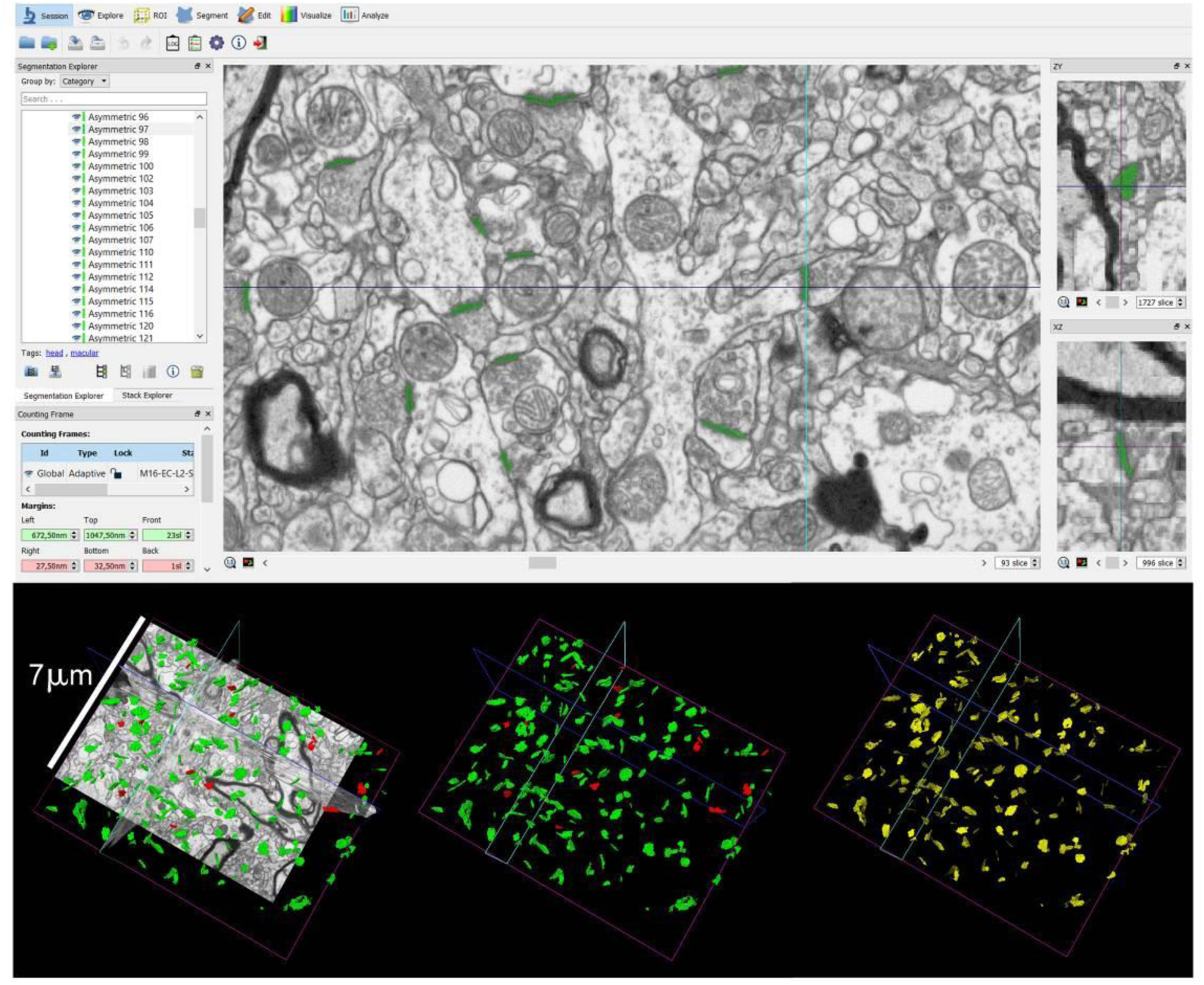
Screenshot of the EspINA software user interface. (Top) In the main window, the sections are viewed through the xy plane (as obtained by FIB/SEM microscopy). The other two orthogonal planes, yz and xz, are also shown in adjacent windows (on the right). (Bottom) The 3D windows show the three orthogonal planes and the 3D reconstruction of AS (green) and SS (red) segmented synapses (bottom left), the reconstructed synapses (bottom center), and the computed SAS for each reconstructed synapse (in yellow; bottom right).

In addition, geometrical features —such as size and shape— and spatial distribution features (centroids) of each reconstructed synapse were also calculated by EspINA. This software also extracts the Synaptic Apposition Area (SAS) and provides its morphological measurements (Fig. 3). Since the pre- and post-synaptic densities are located face to face, their surface areas are comparable (for details, see Morales et al., 2013). Since the SAS comprises both the active zone and the PSD, it is a functionally relevant measure of the size of a synapse (Morales et al., 2013).

To identify the postsynaptic targets of the synapses, we navigated the image stack using EspINA to determine whether the postsynaptic element was a dendritic spine (spine or spines, for simplicity) or a dendritic shaft. Unambiguous identification of spines requires the spine to be visually traced to the parent dendrite. Similarly, for dendritic shafts to be unambiguously identified, they must be visually followed inside the stack. Accordingly, when the postsynaptic element of a synapse was close to the margins and was truncated by the borders of the stack, the identity of the postsynaptic target could not be determined. Therefore, the targets of synapses in each of the stacks were classified into two main categories: spines and dendritic shafts, while truncated elements that could not be safely identified were discarded. When the postsynaptic target was a spine, we further recorded the position of the synapse on the head or neck. We also recorded the presence of single or multiple synapses on a single spine.

### Spatial Distribution Analysis of Synapses

To analyze the spatial distribution of synapses, spatial point-pattern analysis was performed as described elsewhere (Anton-Sanchez et al., 2014; Merchán-Pérez et al., 2014). Briefly, we compared the actual position of centroids of synapses with the Complete Spatial Randomness (CSR) model — a random spatial distribution model which defines a situation where a point is equally likely to occur at any location within a given volume. For each of the 24 different samples, we calculated three functions commonly used for spatial point-pattern analysis: G, F and K functions (for a detailed description, see Blazquez-Llorca et al., 2015). This study was carried out using the Spatstat package and R Project program (Baddeley et al., 2015).

### Statistical analysis

To determine possible differences between layers, statistical comparisons of synaptic density, as well as size of the SAS, were carried out using the unpaired Mann-Whitney (MW) nonparametric U-test (when the normality and homoscedasticity criteria were not met) or t-test parametric test (when the normality and homoscedasticity criteria were met). To identify possible differences within a layer regarding the synaptic size (SAS) related to the shape of the synapses and their postsynaptic target, a Kruskal–Wallis (KW) nonparametric test (when the normality and homoscedasticity criteria were not met) or ANOVA test were performed (when the normality and homoscedasticity criteria were met). Frequency distribution analysis of the SAS was performed using Kolmogorov-Smirnov (KS) nonparametric test. To perform statistical comparisons of AS and SS proportions regarding their synaptic morphology and their postsynaptic target, chi-square (χ^2^) test was used for contingency tables. The same method was used to study whether there were significant differences between layers in relation to the shape of the synaptic junctions and their postsynaptic target.

Statistical studies were performed using the GraphPad Prism statistical package (Prism 7.00 for Windows, GraphPad Software Inc., USA).

## Results

Coronal sections of the human EC at medial level were used for the present study (reviewed in Insausti et al., 2017). The EC was delimited by combining the Nissl and anti-NeuN markers (Fig. 1). The main cytoarchitectural characteristic of layer II (also called Pre-α) is the presence of large islands of modified pyramidal neurons and stellate cells (Braak and Braak, 1992; Insausti and Amaral, 2012; Kobro-Flatmoen and Witter, 2019).

### Synaptic Density

All the synapses reported, counted and analyzed in the present study were individually identified and segmented. The synaptic density values were obtained by dividing the total number of synapses included within the stereological counting frame (CF) by the total volume of the CF.

In the layer II samples, a total of 2,310 synapses were identified and reconstructed in 3D, of which 1,690 synapses were analyzed, after discarding incomplete synapses or those that were touching the exclusion edges of the stereological CF in a total volume of 4,221 μm^3^. Similarly, in the layer III samples, a total of 2,580 synapses were identified and reconstructed in 3D, of which 1,877 synapses were analyzed in a total volume of 4,371 μm^3^.

The average synaptic density of the EC layer II was 0.40 synapses/μm^3^ (with a range of 0.38−0.43 synapses/μm^3^; Table 2; Supplementary figure 1; Supplementary table 1). In layer III, synaptic density was 0.43 synapses/μm^3^ (with a range of 0.41−0.44 synapses/μm^3^; Table 2; Supplementary figure 1; Supplementary table 1). No differences in synaptic density between layers were observed (MW, p=0.18).

### Proportion of excitatory and inhibitory synapses

Since each synapse was fully reconstructed in 3D, it was possible to precisely distinguish between AS and SS (Figs. 2, 3).

In layer II of the EC, the proportion of AS:SS was 92:8, and in layer III this ratio was 93:7 (Table 2; Supplementary table 1). To statistically evaluate whether this difference in the ratio between the two layers was significant, chi-square (χ^2^) tests were applied to 2×2 contingency tables, considering the two layers versus the two types of synapses, and no significant differences were found between layers (χ^2^, p=0.18).

### Three-dimensional spatial synaptic distribution

To analyze the spatial distribution of the synapses, the actual position of each of the synapses in each stack of images was compared with a random spatial distribution model CSR. For this, the functions G, K and F were calculated. In the image stacks analyzed (12 from EC layer II and 12 from EC layer III), the three spatial statistics functions resembled the theoretical curve that simulates the random spatial distribution pattern, which indicated that synapses fitted a random spatial distribution model (Supplementary figure 2).

### Study of the characteristics of the synapse

#### Synaptic size

The study of the synaptic size was carried out analyzing the characteristics of the area and the perimeter of the SAS of each synapse identified and 3D reconstructed in all samples (Fig. 3).

In layer II, the average size (measured by the area of the SAS) of the AS was 110,311 nm^2^, and 65,997 nm^2^ in the case of SS (Table 3; Supplementary table 2). When the average sizes of the two types of synapses were compared, it was found that the area of the AS were significantly larger than the area of the SS (MW, p = 0.03). This difference was also found in the frequency distribution of the area (KS, p <0.0001), indicating that the AS were larger than the SS.

**Table 3.** Area (nm^2^) and perimeter (nm) of the SAS from layer II and layer III of the EC. All data are corrected for shrinkage factor and are expressed as mean± sem. The data for individual cases are shown in Supplementary table 2. AS: asymmetric synapses; EC: entorhinal cortex; SAS: synaptic apposition surface; sem: standard error of the mean; SS: symmetric synapses.

| EC layer | Type of synapse | Area of SAS (nm<sup>2</sup>) | Perimeter of SAS (nm) |
| --- | --- | --- | --- |
| **II** | AS | 110311±3229 | 1631±47 |
|  | SS | 65997±6041 | 1404±73 |
| **III** | AS | 124183±3265 | 1826±67 |
|  | SS | 67445±3243 | 1399±75 |

In layer III, the average area of the AS was 124,183 nm^2^, and in the SS, it was 67,445 nm^2^ (Table 3; Supplementary table 2). When comparing the two types of synapses, it was found that both the area and the perimeter of the AS were significantly larger than the same parameters in the SS (MW, p =0.03). These differences were also found in the frequency distributions of the area and perimeter (KS, p <0.0001), indicating that the AS were larger than the SS, as occurred in layer II.

Statistical comparisons between layers showed differences in the average size of the AS area (MW, p=0.03; Fig. 4A). This statistical difference between layers was also found in the frequency distribution of the area and perimeter of AS (KS, p <0.0001; Fig. 4B, 4C). In summary, AS in layer III were larger than in layer II, while no differences were found for SS.

**Figure 4.**
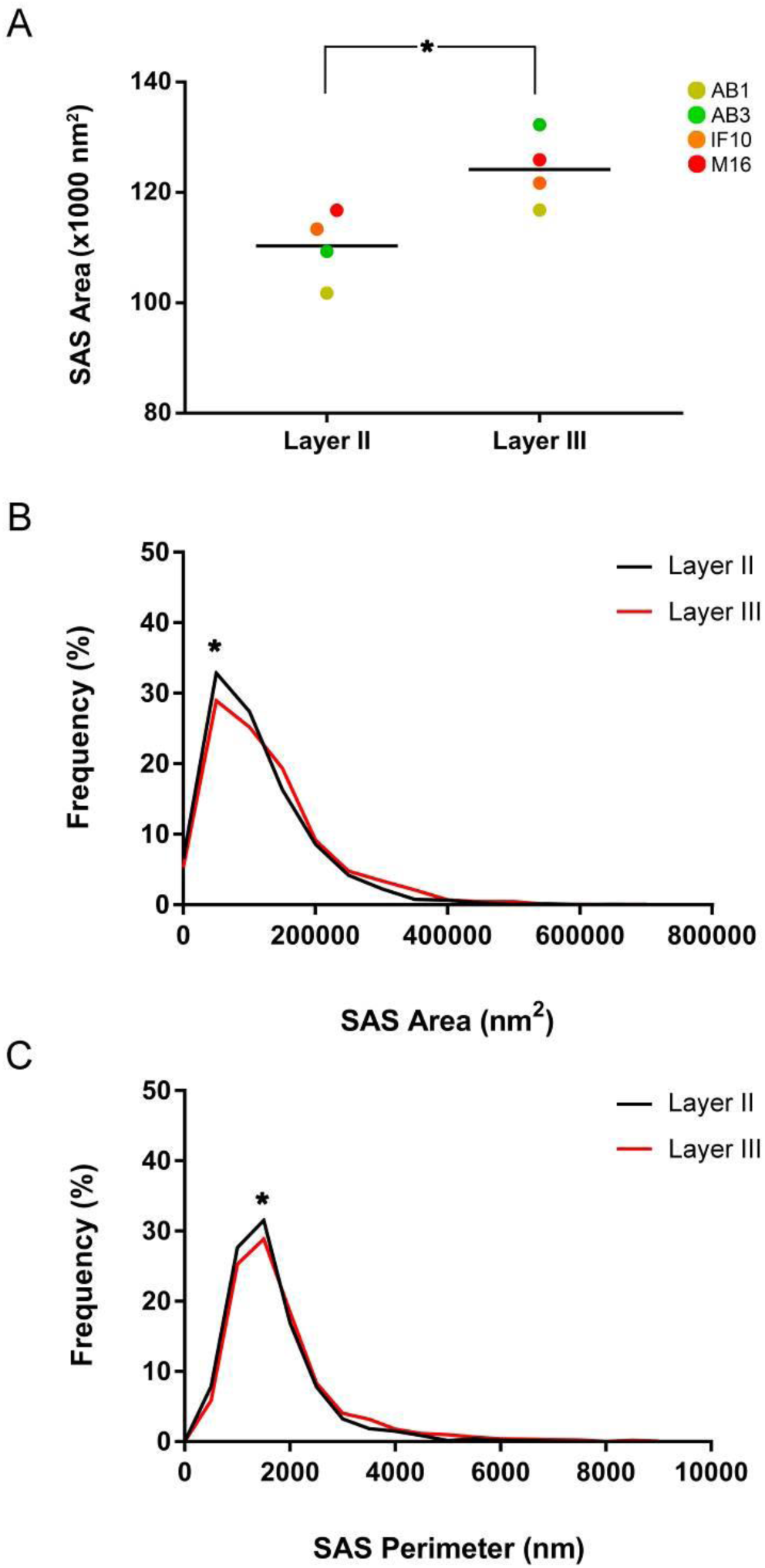
Graph showing the mean AS SAS area (A), and the frequency distribution plots of AS SAS area (B) and perimeter (C), in layers II and III of the EC. (**A**) A different color corresponds to each analyzed case, as denoted in the key. Statistical comparisons between layers showed differences in the size of the AS area (**B**) (asterisk; MW, p=0.03), as well as in the frequency distribution of the perimeter (**C**) (KS, p<0.0001; indicated with an asterisk).

#### Synaptic shape

Regarding shape, the synapses were classified into four categories: macular (with a flat, disk-shaped PSD), perforated (with one or more holes in the PSD), horseshoe (with an indentation in the perimeter of the PSD) or fragmented (with two or more physically discontinuous PSDs) (Supplementary Fig. 3A; for a detailed description, see Domínguez-Álvaro et al., 2019; Santuy et al., 2018a).

In layer II, a total of 1,549 AS were identified and reconstructed in 3D, with 83% presenting a macular morphology, followed by 12.9% perforated, 3.3% horseshoe and 0.8% fragmented (Fig. 5A). As for the SS, a total of 137 synapses were identified and reconstructed in 3D, with the majority presenting a macular morphology (73%). Of the remaining SS, 15.3% were horseshoe-shaped, 11% were perforated and 0.7% were fragmented (Table 4; Supplementary table 3). Determining the proportions of the two categories (i.e., AS and SS; Fig. 5B) for each synapse shape revealed that, of the total macular synapses, 92.8% were AS and 7.2% were SS. This proportion was maintained in the case of perforated synapses (93% versus 7% SS), while it changed significantly in the case of the horseshoe synapses, where 70.8% were AS and 29.2% were SS (χ^2^, p <0.0001). Thus, SS showing a horseshoe shape were more frequent than expected according to the general proportion of SS (7%). In the case of fragmented synapses, the proportion was maintained (92.9% AS and 7.1% SS).

**Table 4.** Proportion of the different shapes of synaptic junctions in layer II and layer III of the EC. Data are given as percentages with the absolute number of synapses studied in parentheses. Data for each individual case are shown in Supplementary table 3. AS: asymmetric synapses; EC: entorhinal cortex; SS: symmetric synapses.

| EC layer | Type of synapse | Macular synapses | Perforated synapses | Horseshoe-shaped synapses | Fragmented synapses | Total synapses |
| --- | --- | --- | --- | --- | --- | --- |
| **II** | AS | 83.0% | 12.9% | 3.3% | 0.8% | 100% |
|  |  | (1285) | (200) | (51) | (13) | (1549) |
|  | SS | 73.0% | 11.0% | 15.3% | 0.7% | 100% |
|  |  | (100) | (15) | (21) | (1) | (137) |
| **III** | AS | 75.9% | 19.9% | 3.3% | 0.9% | 100% |
|  |  | (1326) | (347) | (58) | (15) | (1746) |
|  | SS | 65.9% | 15.5% | 17.0% | 1.6% | 100% |
|  |  | (85) | (20) | (22) | (2) | (129) |

**Figure 5.**
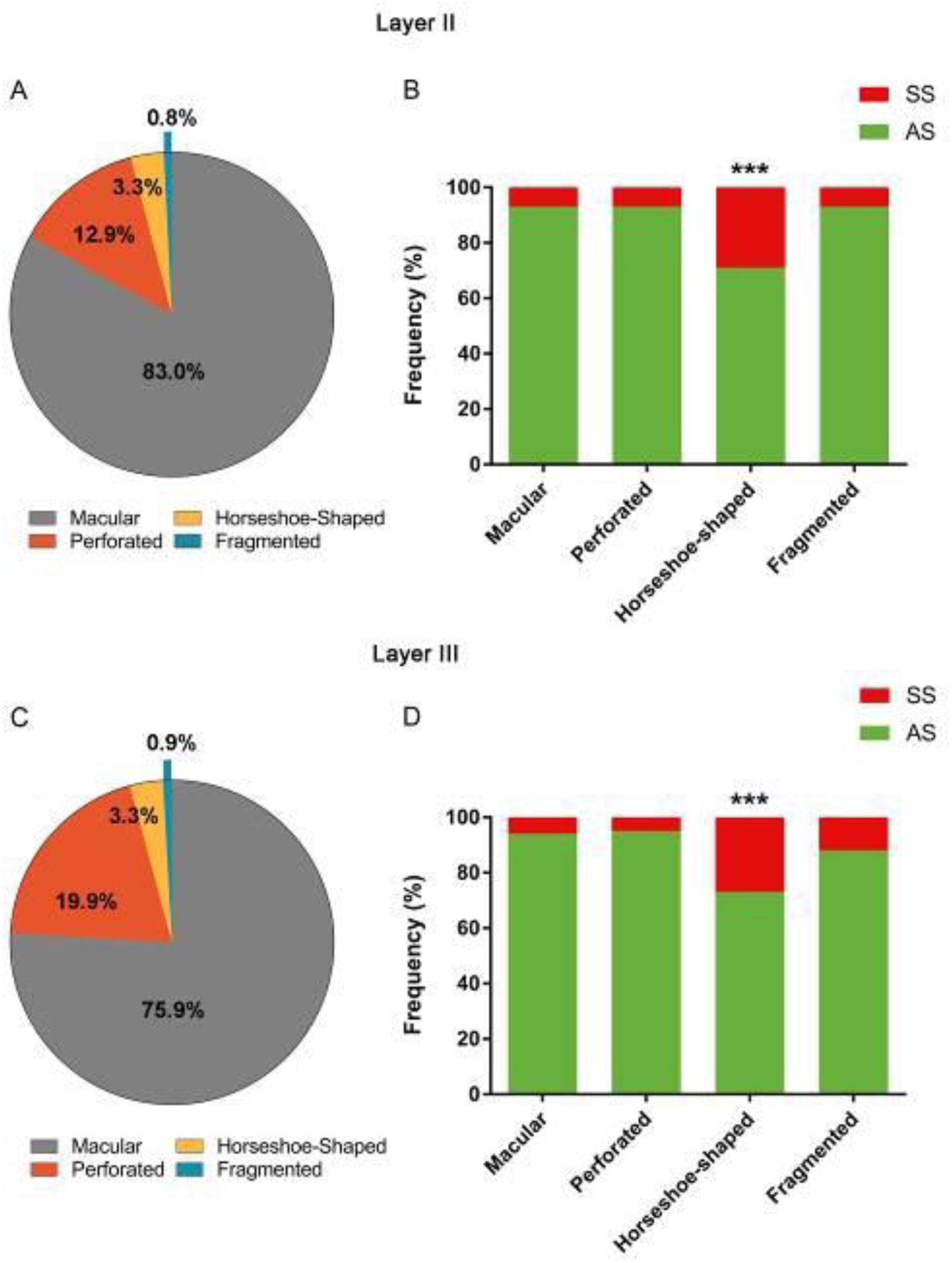
Proportions of the different synaptic shapes in layers II (A, B) and III (C, D) of the EC. (**A)** Shape proportions of the AS in layer II. (**B)** Proportions of AS and SS belonging to each shape type in layer II. (**C)** Shape proportions of the AS in layer III. A significantly fewer number of macular AS and a significantly higher number of perforated AS were found compared to layer II (χ^2^, p <0.0001). (**D)** Proportions of AS and SS belonging to each shape type in layer III. In both layer II and layer III, the horseshoe-shaped synapses were significantly more frequent among SS (*X*^2^, *p*<0.0001).

In the layer III samples, a total of 1,746 AS were identified and reconstructed in 3D. Of these, the majority presented a macular morphology (75.9%), followed by perforated (19.9%) and horseshoe (3.3%), and only a small percentage of the AS presented a fragmented morphology (0.9%) (Fig. 5C). A significantly lower number of macular AS and a significantly higher number of perforated AS were found compared to layer II (χ^2^, p <0.0001). Of a total of 129 SS, 65.9% had a macular morphology, 15.5% had a perforated shape, 17% were horseshoe-shaped, and 1.6% had a fragmented morphology (Table 4; Supplementary table 3). Regarding the prevalence of each type (AS and SS (Fig. 5D)), 94% of the macular morphology synapses were AS, while 6% were SS. This proportion was maintained in the perforated synapses (94.5% AS versus 5.5% SS); however, similar to the observations in layer II, this changed significantly in the case of horseshoe-shaped synapses, where 72.5% were AS and 27.5% were SS (χ^2^, p <0.0001). Thus, SS showing a horseshoe shape were more frequent than expected according to the general proportion of SS. In the case of the fragmented synapses, 88.2% corresponded to AS, while 11.8% were SS.

#### Synaptic Size and Shape

We also determined whether the shape of the synapses was related to their size. For this purpose, the area and perimeter of the SAS, of both AS and SS, were analyzed according to synaptic shape.

We found that, in both layer II and III samples, the area and perimeter of the macular AS were significantly smaller than the area and the perimeter of the perforated, horseshoe or fragmented AS (ANOVA, p <0.001; Supplementary figure 3B–E). This tendency was also observed in SS (but no significant differences were found).

Only differences were found in the synaptic size (measured as the average of the AS area and perimeter) of the fragmented synapses between the two layers (ANOVA, p <0.0001). However, the number of fragmented synapses was not large enough to draw statistically reliable conclusions. No differences were found in the frequency distribution of the area and perimeter of AS (KS, p >0.01).

### Study of the postsynaptic elements

Postsynaptic targets were unambiguously identified and classified as spines and dendritic shafts. Additionally, when the postsynaptic element was a spine, we distinguished the location of the synapse on the neck or on the head of this spine. When the postsynaptic element was identified as a dendritic shaft, it was classified as “with spines” or “without spines” (for details, see Domínguez-Álvaro et al., 2019).

The postsynaptic elements of 1,157 AS and 124 SS were determined in the layer II samples — 50.3% of the AS were established on dendritic shafts (25.8% on dendritic shafts with spines and 24.5% on dendritic shafts without spines), 49.0% of the AS were established on spine heads and 0.7% on spine necks. In the case of SS, 88% were established on dendritic shafts (63% on dendritic shafts with spines and 25% on dendritic shafts without spines), while 10.4% were established on spine heads, and the remaining 1.6% were on spine necks (Table 5; Supplementary table 4).

**Table 5.** Distribution of AS and SS on spines and dendritic shafts in layer II and layer III of the EC. Synapses on spines have been sub-divided into those established on spine heads and those established on spine necks. Moreover, we differentiated between aspiny and spiny dendritic shafts. Data are given as percentages with the absolute number of synapses studied in parentheses. Data for each individual case are shown in Supplementary table 4. AS: asymmetric synapses; EC: entorhinal cortex; SS: symmetric synapses.

| EC layer | Type of synapse | Synapses on spine heads | Synapses on spine necks | Synapses on aspiny dendritic shaft | Synapses on spiny dendritic shaft | Total synapses |
| --- | --- | --- | --- | --- | --- | --- |
| **II** | AS | 49.0% | 0.7% | 24.5% | 25.8% | 100% |
|  |  | (567) | (8) | (284) | (298) | (1157) |
|  | SS | 10.4% | 1.6% | 25.0% | 63.0% | 100% |
|  |  | (13) | (2) | (31) | (78) | (124) |
| **III** | AS | 46.7% | 0.9% | 20.6% | 31.8% | 100% |
|  |  | (639) | (12) | (281) | (435) | (1367) |
|  | SS | 13.3% | 2.5% | 22.5% | 61.7% | 100% |
|  |  | (16) | (3) | (27) | (74) | (120) |

When the preference of the AS and the SS for a specific postsynaptic element was analyzed, it was found that the AS showed a statistically significant preference for spine heads (χ^2^, p <0.0001), while the SS showed a statistically significant preference for dendritic shafts (χ^2^, p <0.0001; Fig. 6A). Considering all types of synapses established on the spine heads, the proportion of AS:SS was 98:2; while in those established on dendritic shafts, this proportion was 84:16. Since the overall AS:SS ratio in layer II was 92:8, the present results show that AS and SS did show a preference for a particular postsynaptic element, that is, the AS showed a preference for the spine heads, while the SS showed a preference for the dendritic shafts.

**Figure 6.**
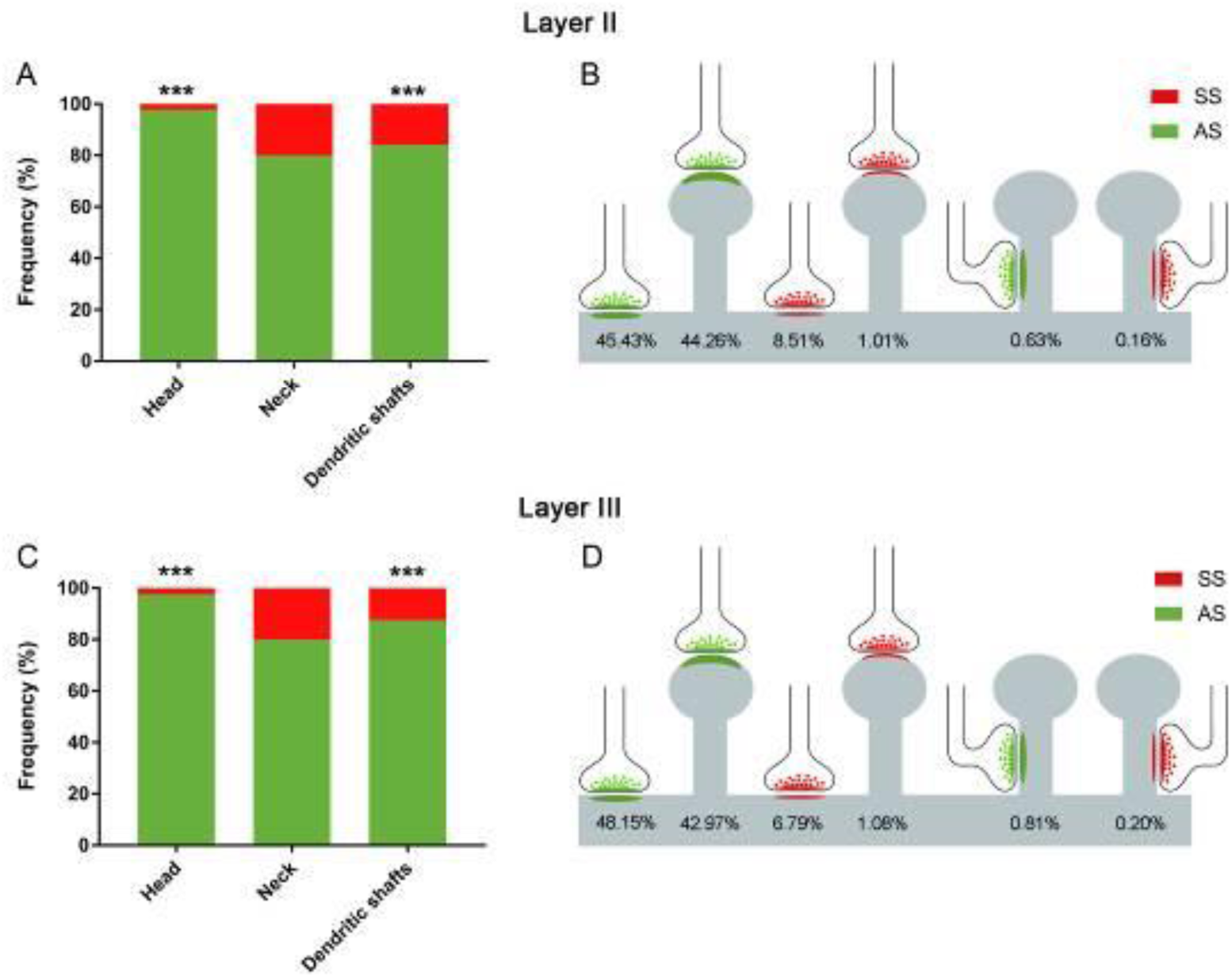
Analysis of the postsynaptic target distribution in layers II (A, B) and III (C, D) of the EC. **(A)** Graph showing the proportions of AS and SS corresponding to each postsynaptic target in layer II. (**B)** Schematic representation of the distribution of AS and SS on different postsynaptic targets in layers II. **(C)** Proportions of AS and SS corresponding to each postsynaptic target in layer III. In both, layer II and layer III, the AS showed a preference for spine heads, while the SS showed a preference for dendritic shafts (asterisks; χ^2^, p<0.0001). (**D)** Schematic representation of the distribution of AS and SS on different postsynaptic targets in layer III. (**B, D)** Percentages of postsynaptic targets are indicated, showing —from left to right— the most frequent type (AS on dendritic shafts) to the least frequent type (SS on spine necks). (**A−D**) Synapses on spines have been sub-classified into those that are established on the spine head and those established on the neck.

Grouping by both type of synapse (AS or SS) and type of postsynaptic target (spine head, spine neck or dendritic shaft), it was found that 45.4% of layer II synapses were AS established on dendritic shafts, closely followed by AS on spine heads (44.3%), while 8.5% were SS on dendritic shafts and 1.0% were SS on spine heads. The two least frequent types of synapses were AS and SS established on spine necks (0.6% and 0.2%, respectively; Fig. 6B).

Regarding the samples of layer III, the postsynaptic elements of 1,367 AS and 120 SS were analyzed; 52.4% of the AS were located on dendritic shafts (20.6% on dendritic shafts without spines, 31.8% on dendritic shafts with spines), while 46.7% of AS were established on spine heads and 0.9% on necks. Most of the SS were established on dendritic shafts (61.7% on dendritic shafts with spines and 22.5% on dendritic shafts without spines), while 13.3% were found on spine heads, and 2.5% on necks (Table 5; supplementary table 4).

When the preference of the synaptic types (AS or SS) for a particular postsynaptic element was analyzed, we found that the AS presented a statistically significant preference for spine heads (χ^2^, p <0.0001); while the SS showed a significant preference for dendritic shafts (χ^2^, p <0.0001; Fig. 6C), in line with what we found in layer II. Analysis of synapses established on spine heads showed an AS:SS ratio of 98:2, whereas —in dendritic shafts— the AS:SS ratio was 88:12. Given that the general ratio of AS and SS in layer III was 92:8, it could be concluded that the AS presented preference for the spine heads and the SS for the dendritic shafts, similar to our observations in layer II. In the case of synapses on dendritic necks, although a higher proportion of SS was observed compared to what would be expected by chance, the SS sample was not large enough to draw statistically reliable conclusions.

Simultaneously considering synaptic types and postsynaptic targets, we found similar results to layer II: 48.1% were AS on dendritic shafts, followed by AS on spine heads (43.0%), while 6.8% were SS on dendritic shafts and 1.1% were SS on spine heads. The less frequent types of synapses were AS and SS established on spine necks (0.8% and 0.2%, respectively (Fig. 6D)).

Statistical comparisons of the proportion of AS (χ^2^, p=0.30) and SS (χ^2^, p=0.46) established on spines and dendritic shafts did not show differences between layers II and III.

Finally, to detect the presence of multiple synapses, an analysis of the synapses established on spine heads was performed. In layer II, the most frequent finding was a single AS per head (90.3%), followed by two AS (5.5%), while 3.8% had one AS and one SS on a head, and the frequency of a single SS per head was very low (0.4%; Fig. 7). Likewise, in layer III, the most frequent finding was the presence of a single AS on a head (89.6%), followed by two AS (6.1 %), while only 3.7% had one AS and one SS on the same head, and the least frequent combinations were a single SS and two SS on a head (0.3% and 0.3%, respectively; Fig. 7).

**Figure 7.**
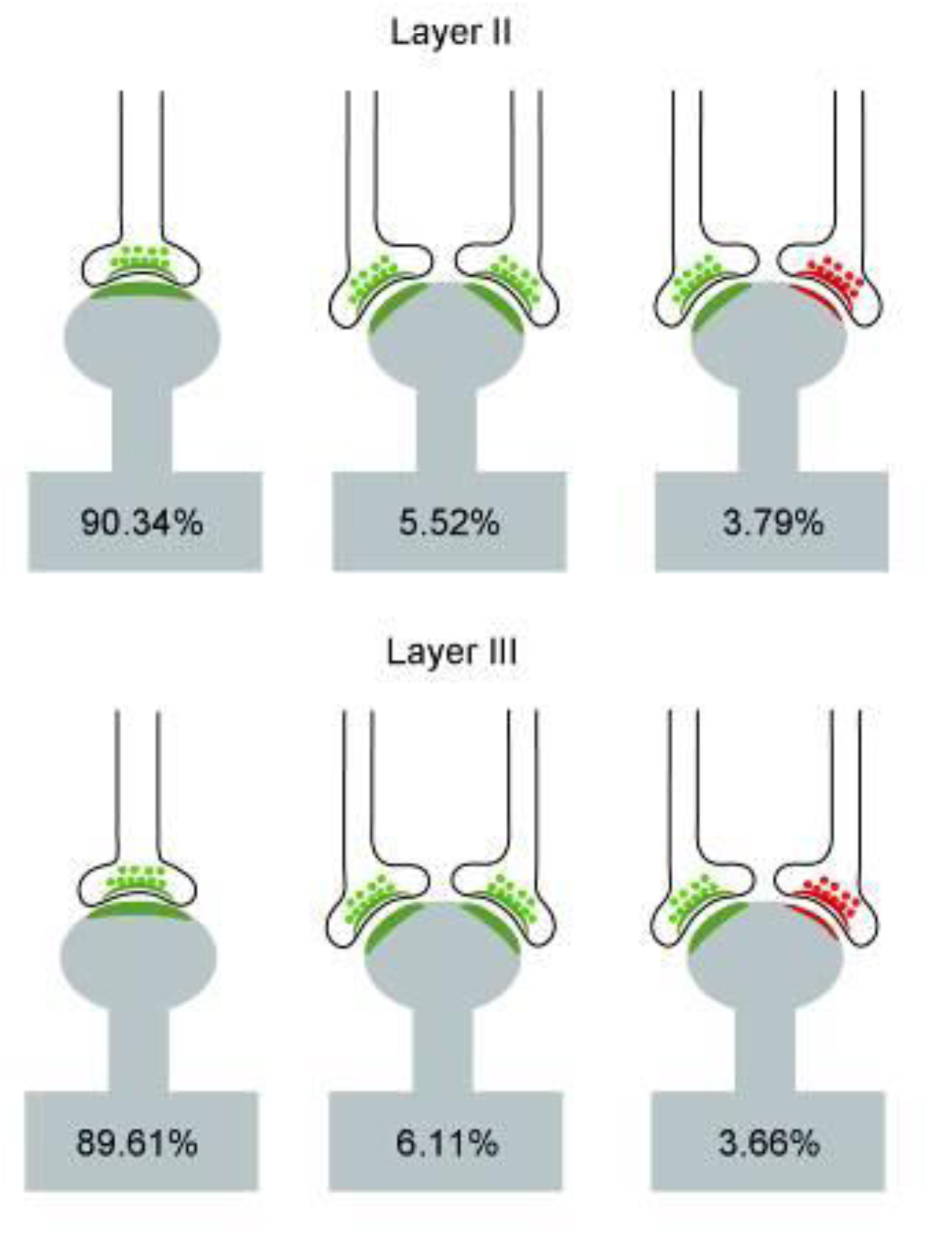
Schematic representation of single and multiple synapses on dendritic spine heads in layers II and III of the EC. Percentages of each type are indicated. Synapses on the necks and other combinations that were rarely found (less than 1%) have not been included. AS have been represented in green and SS in red.

#### Postsynaptic elements and synaptic size

In addition to the study of the distribution of synapses in the different postsynaptic elements, we analyzed whether there was a relationship between the synapse size and the type of postsynaptic element. The study was carried out with the data of the area and perimeter of the SAS of each synapse whose postsynaptic element was identified.

In layer II, AS established on spine necks were smaller than AS established on dendritic heads or shafts (ANOVA, p <0.0001; Supplementary figure 4). However, in the samples from layer III, no statistically significant differences were found in the size of the AS established on spine heads, spine necks or dendritic shafts (ANOVA, p> 0.05; Supplementary figure 4). In the case of SS, their numbers were too low to perform a robust statistical analysis.

The comparison between layers regarding the size of the synapses according to the postsynaptic element did not show any difference (ANOVA, p> 0.05). However, we found differences between layers regarding the frequency distribution of the SAS perimeter of the AS established on dendritic shafts (KS, p=0.0005) — larger synapses were more frequent on the dendritic shafts in layer III. This finding is in line with the results in layer III regarding the larger size of AS and the higher proportion of perforated AS synapses.

## Discussion

The present study constitutes a detailed description of the synaptic organization of the neuropil of layers II and III from the human EC using 3D EM. As far as we know, detailed 3D synaptic morphology analysis and identification of postsynaptic targets at the ultrastructural level have not been performed before in the human EC. Determination of the postsynaptic targets, as well as the shape and size of the synaptic junctions, provides critical data on synaptic functionality. The present results have yielded a new, large, quantitative ultrastructure dataset in 3D of the synaptic organization of this particular brain region.

At the ultrastructural level, the following major results were obtained: (i) synaptic density and ratio were quite similar in both layers; (ii) synapses fitted into a random spatial distribution in both layers; (iii) in both layers, AS were larger than SS, and in layer III, AS were larger than in layer II; (iv) regardless of the layer, most synapses are AS and display a macular shape — and there is an almost equal split between the targeting of dendritic shafts and spines.

### Synaptic density

Synaptic density is critical not only to describe the synaptology of a particular brain region, but also in terms of connectivity. The mean synaptic density in EC was 0.42 synapses/µm^3^ and was similar in layer II (0.40 synapses/µm^3^) and III (0.43 synapses/µm^3^). These values are higher than those reported by Scheff et al. (1993) using transmission electron microscopy (TEM). These authors estimated a density of 0.32 synapses/µm^3^ in layer III of EC. This difference between our results and those of Scheff et al. may arise not only from differences in age of the human brain tissue used —4 individuals aged 40, 45, 53 and 65 years old in our study versus 11 individuals aged 60−83 years old in the study by Scheff et al. (1993)—, but also from the method used to estimate synaptic density. Perhaps more importantly, in the TEM study, synaptic density was calculated using estimations based on the number of synaptic profiles per unit area (size-frequency method) in single ultrathin sections of tissue. That is, the number of synapses per volume unit is estimated from two-dimensional samples. As previously discussed (Merchán-Pérez et al., 2009), FIB/SEM technology provides the *actual* number of synapses per volume unit, and it avoids most of the errors associated with stereological methods. In addition, our current data are very similar within cases, showing very little variability between samples (Supplementary table 1; Supplementary figure 1), suggesting a robust estimation of the synaptic density using FIB/SEM in these EC layers.

Our previous 3D ultrastructural study using FIB/SEM in layer II of the human transentorhinal cortex (TEC) showed a mean synaptic density of 0.41, 0.42 and 0.43 synapses/µm^3^ in cases AB1, IF10 and M16, respectively — three cases that were also analyzed in the present study (Domínguez-Álvaro et al., 2018). Moreover, estimations performed in human CA1 hippocampal region, also using FIB/SEM technology, have shown (in cases AB1 and AB3) a synaptic density in the stratum oriens ranging from and 0.43 synapses/µm^3^, respectively, and 1.09 synapses/µm^3^ in the superficial part of the stratum pyramidale in both cases (Montero-Crespo et al., 2020). Since the processing and analysis methods were identical, these similarities (with layer II from TEC and stratum oriens of CA1) and differences (with CA1 pyramidal layer) may be attributable to the specific layer and brain region analyzed, suggesting that synaptic density greatly depends on the human brain layer and/or region being studied.

### Proportion of synapses and spatial synaptic distribution

At the circuit level, the proportion of excitatory and inhibitory synapses is critical from a functional point of view, since higher or lower proportions linked to differences in the excitatory/inhibitory balance of the cortical circuits (for reviews, see Froemke, 2015; Zhou and Yu, 2018; Sohal and Rubenstein, 2019). It is well established, in different brain regions and species, that the cortical neuropil has many more excitatory synapses than inhibitory synapses (reviewed in DeFelipe et al., 2002; DeFelipe, 2011). In the present study, the proportion of excitatory synapses was higher in layers II and III (the AS:SS ratios were 92:8 and 93:7, respectively). In general, similar results have been found in other brain regions using the same method. Our previous studies have shown relatively small differences: an AS:SS ratio of 96:4 in the TEC (Domínguez-Álvaro et al., 2018), and 95:5 in the CA1 hippocampal field (Montero-Crespo et al., 2020).

Regarding the spatial organization of synapses, we found that synapses were randomly distributed in the neuropil of both layers. This type of spatial distribution has also been found in the frontal and transentorhinal cortices, as well as in the CA1 hippocampal region of the human brain (Blazquez-Llorca et al., 2013; Domínguez-Álvaro et al., 2018; Montero-Crespo et al., 2020).

Therefore, the present results suggest that the random spatial synaptic distribution of synapses and the AS:SS ratio (ranging from 92 to 96% for AS and 8 to 4% for SS) are widespread ‘rules’ of the synaptic organization of the human cerebral cortex.

### Shape and size of the synapses

There are very few studies on human brain that provide data regarding the size of the synaptic junctions for comparison with our current results. Similar studies have been performed in layer IV of the temporal neocortex (Yakoubi et al., 2019). Yakoubi’s group analyzed 150 boutons establishing asymmetric synapses and reported that the mean size of the active zone was 130,000 nm^2^. These results are in line with our current measurements for the area of the SAS of AS in both layer II (110,311 nm^2^) and layer III (124,183 nm^2^), and also similar to our previous results in the human TEC (118,037 nm^2^; Domínguez-Álvaro et al., 2018). In our studies on human EC and TEC, 3,300 and 2,545 axon terminals forming AS were analyzed, respectively, and they therefore represent robust data.

We also found that, in both EC layers, excitatory contacts (AS) were larger than inhibitory (SS) ones (117,247 vs. 66,721 nm^2^, respectively), as was also observed in layer II of the human TEC (118,037 vs. 73,590 nm^2^; Domínguez-Álvaro et al., 2019). Furthermore, we found that AS of layer III were significantly larger (124,183 nm^2^) than those in layer II (110,311 nm^2^), suggesting layer specific characteristics regarding synaptic size. Synaptic size has been proposed to correlate with release probability, synaptic strength, efficacy and plasticity (Nusser et al., 1998; Takumi et al., 1999; Ganeshina et al., 2004a; Tarusawa et al., 2009; Südhof, 2012; Holderith et al., 2012; Biederer et al., 2017).

Most EC synapses presented a macular shape (taking both layers together, 78% were macular), whereas 21% were the more complex-shaped synapses (perforated, horseshoe and fragmented) — both figures that are comparable to previous reports in different brain areas and species (Domínguez-Álvaro et al., 2019; Montero-Crespo et al., 2020; Geinisman et al., 1987; Jones and Calverley, 1991; Neuman et al., 2015; Santuy et al., 2018b). Complex-shaped synapses are larger than macular ones. In particular, perforated synapses have more AMPA and NMDA receptors than macular synapses and are thought to constitute a relatively ‘powerful’ population of synapses with more long-lasting memory-related functionality than macular synapses (Ganeshina et al., 2004a, b; Spruston, 2008; Vicent-Lamarre et al., 2018). Our current results also showed that macular AS were smaller than the complex-shaped synapses. In addition, macular synapses were less abundant in layer III than in layer II. Considering all types of synapses, AS in layer III were larger than those in layer II.

As mentioned above, neurons in layer III of the EC project to CA1 and the subiculum, whereas those in layer II send their axons to DG and CA3. Moreover, upper EC layers receive cortical afferents from a number of regions as well as from the parahippocampal and perirhinal cortex (Schultz et al., 2015). Therefore, the differences in synaptic size observed between layer II and III may reveal unique microanatomical synaptic features that may be related to the differential pattern of layer connectivity in human EC.

### Postsynaptic targets

A clear preference of excitatory axons for dendritic spines and inhibitory axons for dendritic shafts was observed, which has also been found in a large variety of cortical regions and species, including humans (reviewed in DeFelipe et al., 2002), although very few studies have been performed in the human brain.

In the present work, the co-analysis of the synaptic type and postsynaptic target showed that the proportion of AS established on dendritic spines (also known as ‘axospinous’) reached around 45−48% in both layer II and layer III of the EC. Using the same FIB/SEM technology, in layer II of the human TEC, we found that 55% of AS were established on dendritic spines (Domínguez-Álvaro et al., 2019), whereas in the superficial part of the CA1 stratum pyramidale, this percentage reached 93%, and the other CA1 sublayers examined also displayed high proportions of axospinous AS (stratum oriens: 84%, deep stratum pyramidale: 90%, stratum radiatum: 87%, stratum lacunosum-moleculare: 72%; Montero-Crespo et al., 2020). Furthermore, serial electron microscopy studies performed in layer IV and V of the human temporal neocortex have reported that axospinous AS account for approximately 77% and 85% of the AS, respectively (Yakoubi et al., 2019a, b), whereas our preliminary data from FIB/SEM studies in layer III of the human temporal cortex —based on the 3D reconstruction of 1,456 synapses— showed that 68% of AS were axospinous (Cano-Astorga et al., in preparation). Finally, there are important differences and similarities in the proportion of AS on spines and dendritic shafts in different cortical regions and layers, which may represent another microanatomical specialization of the cortical regions examined.

Although our data is very robust since it is based on the analysis of thousands of synapses, the data was obtained from four individuals. Therefore, caution must be exercised when interpreting the significance of the results. Performing further studies on brain tissue from more subjects of different ages — including both males and females

— will be necessary to better understand both the variability and invariance of human synaptic organization.

In conclusion, the present work constitutes a detailed description of the synaptic organization of the human EC, which is a necessary step to better understand its functional organization in both health and disease.

## Abbreviation list

3D: three-dimensional,
AS: asymmetric synapses,
CA1: *cornu ammonis* 1,
CSR: Complete Spatial Randomness,
DG: dental gyrus,
EC: entorhinal cortex,
FIB/SEM: focused ion beam/scanning electron microscopy,
KS: Kolmogorov-Smirnov,
MW: Mann-Whitney,
PB: phosphate buffer,
SAS: synaptic apposition surface,
SD: standard deviation,
sem: standard error of the mean,
SS: symmetric synapses

## Ethics approval

Brain tissue samples were obtained following the guidelines and approval of the Institutional Ethical Committee from all involved institutions: Bellvitge University Hospital (Barcelona, Spain); *Centro Alzheimer Fundación Reina Sofía*, CIEN Foundation (Madrid, Spain); and School of Medicine, University of Castilla-la Mancha, (Albacete, Spain).

## Availability of data and materials

The datasets used and analyzed during the current study are available within the article or its supplementary materials, and from the corresponding author on reasonable request.

## Authors’ contributions

JDeF and LA-N oversaw and designed the project. MD-A and LA-N designed and performed experiments. MD-A performed data analysis. LB-L and MM-C performed experiments and helped interpret experiments. LA-N drafted the initial manuscript. All authors read, reviewed, edited, and approved the final manuscript.

## Acknowledgments

We would like to thank Carmen Álvarez and Lorena Valdés for their technical assistance, and Nick Guthrie for his excellent text editing.

---

**Supplementary table 1 (for Table 2).**
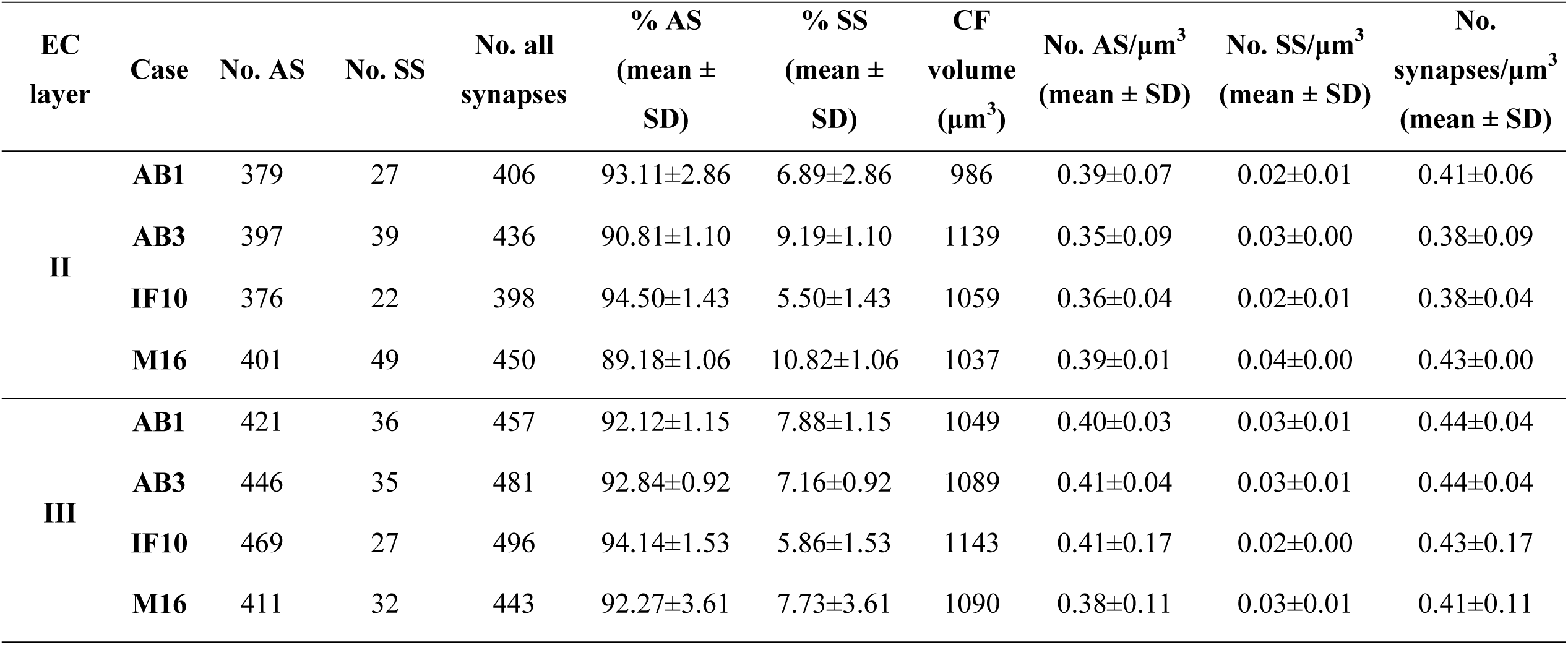
Accumulated data obtained from the ultrastructural analysis of neuropil from layers II and III of the EC by individual cases. All volume data are corrected for shrinkage factor. AS: asymmetric synapses; CF: counting frame; EC: entorhinal cortex; SD: standard deviation; SS: symmetric synapses.

| EC layer | Case | No. AS | No. SS | No. all synapses | % AS<br>(mean ± SD) | % SS<br>(mean ± SD) | CF volume<br>($\mu\text{m}^3$) | No. AS/$\mu\text{m}^3$<br>(mean ± SD) | No. SS/$\mu\text{m}^3$<br>(mean ± SD) | No. synapses/$\mu\text{m}^3$<br>(mean ± SD) |
| --- | --- | --- | --- | --- | --- | --- | --- | --- | --- | --- |
| II | AB1 | 379 | 27 | 406 | 93.11±2.86 | 6.89±2.86 | 986 | 0.39±0.07 | 0.02±0.01 | 0.41±0.06 |
| | AB3 | 397 | 39 | 436 | 90.81±1.10 | 9.19±1.10 | 1139 | 0.35±0.09 | 0.03±0.00 | 0.38±0.09 |
| | IF10 | 376 | 22 | 398 | 94.50±1.43 | 5.50±1.43 | 1059 | 0.36±0.04 | 0.02±0.01 | 0.38±0.04 |
| | M16 | 401 | 49 | 450 | 89.18±1.06 | 10.82±1.06 | 1037 | 0.39±0.01 | 0.04±0.00 | 0.43±0.00 |
| III | AB1 | 421 | 36 | 457 | 92.12±1.15 | 7.88±1.15 | 1049 | 0.40±0.03 | 0.03±0.01 | 0.44±0.04 |
| | AB3 | 446 | 35 | 481 | 92.84±0.92 | 7.16±0.92 | 1089 | 0.41±0.04 | 0.03±0.01 | 0.44±0.04 |
| | IF10 | 469 | 27 | 496 | 94.14±1.53 | 5.86±1.53 | 1143 | 0.41±0.17 | 0.02±0.00 | 0.43±0.17 |
| | M16 | 411 | 32 | 443 | 92.27±3.61 | 7.73±3.61 | 1090 | 0.38±0.11 | 0.03±0.01 | 0.41±0.11 |

**Supplementary table 2 (for Table 3).** Data regarding area and perimeter of the SAS in layers II and III of the EC by individual cases. All data are corrected for shrinkage factor and are expressed as mean±sem. AS: asymmetric synapses; SS: symmetric synapses.

| EC layer | Case | Type of synapse | Area of SAS (nm<sup>2</sup>) | Perimeter of SAS (nm) |
| --- | --- | --- | --- | --- |
| II | AB1 | AS | 101766±3757 | 1551±40 |
|  |  | SS | 58790±6865 | 1252±117 |
|  | AB3 | AS | 109320±3694 | 1605±38 |
|  |  | SS | 71082±13154 | 1593±226 |
|  | IF10 | AS | 113399±4343 | 1770±51 |
|  |  | SS | 53681±4642 | 1331±112 |
|  | M16 | AS | 116757±4139 | 1598±40 |
|  |  | SS | 80435±6621 | 1440±73 |
| III | AB1 | AS | 116843±3974 | 1673±43 |
|  |  | SS | 60978±5015 | 1220±79 |
|  | AB3 | AS | 132252±4051 | 1859±47 |
|  |  | SS | 67083±5083 | 1444±89 |
|  | IF10 | AS | 121736±4345 | 1991±60 |
|  |  | SS | 65338±9444 | 1578±210 |
|  | M16 | AS | 125900±4790 | 1779±55 |
|  |  | SS | 76379±10422 | 1354±129 |

**Supplementary table 3 (for Table 4).** Proportions of the different shapes of synaptic junctions in layers II and III of the EC by individual cases. Data are given as percentages with the absolute number of synapses studied in parentheses. AS: asymmetric synapses; EC: entorhinal cortex; SS: symmetric synapses.

| EC layer | Case | Type of synapse | Macular synapses | Perforated synapses | Horseshoe-shaped synapses | Fragmented synapses | Total synapses |
| --- | --- | --- | --- | --- | --- | --- | --- |
| II | AB1 | AS | 84.7% (320) | 9.8% (37) | 4.5% (17) | 1.0% (4) | 100% (378) |
|  |  | SS | 74.1% (20) | 7.4% (2) | 18.5% (5) | 0% (0) | 100% (27) |
|  | AB3 | AS | 82.6% (327) | 13.6% (54) | 2.5% (10) | 1.3% (5) | 100% (396) |
|  |  | SS | 66.6% (26) | 15.4% (6) | 15.4% (6) | 2.6% (1) | 100% (39) |
|  | IF10 | AS | 81.6% (307) | 14.1% (53) | 4.0% (15) | 0.3% (1) | 100% (376) |
|  |  | SS | 72.7% (16) | 9.1% (2) | 18.2% (4) | 0% (0) | 100% (22) |
|  | M16 | AS | 83.0% (331) | 14.0% (56) | 2.3% (9) | 0.7% (3) | 100% (399) |
|  |  | SS | 77.6% (38) | 10.2% (5) | 12.2% (6) | 0% (0) | 100% (49) |
| III | AB1 | AS | 78.6% (330) | 17.6% (74) | 3.6% (15) | 0.2% (1) | 100% (420) |
|  |  | SS | 80.0% (28) | 5.7% (2) | 11.4% (4) | 2.9% (1) | 100% (35) |
|  | AB3 | AS | 73.5% (328) | 21.1% (94) | 3.4% (15) | 2.0% (9) | 100% (446) |
|  |  | SS | 48.6% (17) | 14.3% (5) | 37.1% (13) | 0% (0) | 100% (35) |
|  | IF10 | AS | 76.3% (358) | 19.6% (92) | 3.7% (17) | 0.4% (2) | 100% (469) |
|  |  | SS | 66.7% (18) | 22.2% (6) | 11.1% (3) | 0% (0) | 100% (27) |
|  | M16 | AS | 75.4% (310) | 21.2% (87) | 2.7% (11) | 0.7% (3) | 100% (411) |
|  |  | SS | 68.8% (22) | 21.9% (7) | 6.2% (2) | 3.1% (1) | 100% (32) |

**Supplementary table 4 (Table 5).**
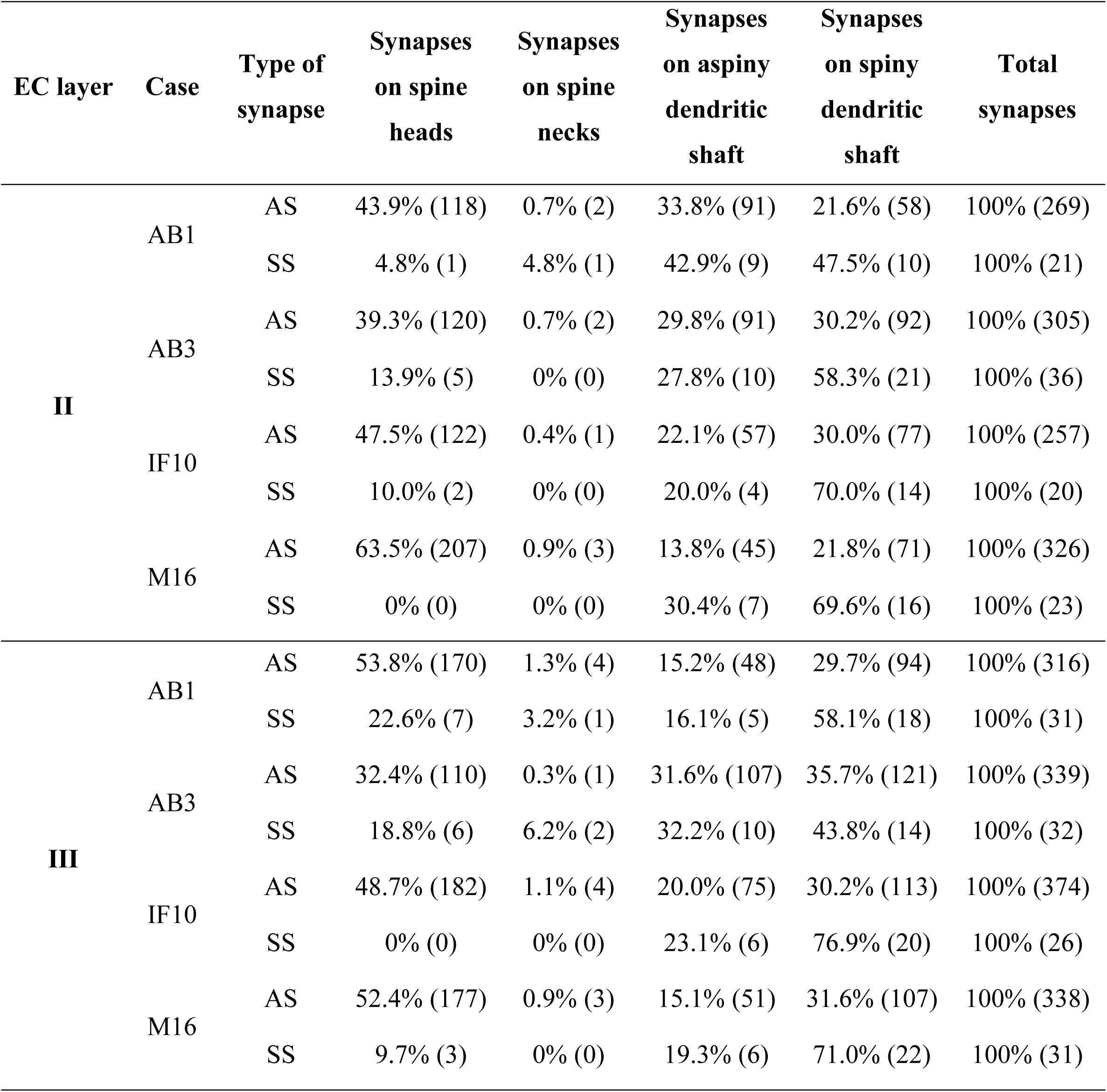
Distribution of AS and SS on spines and dendritic shafts in layers II and III of the EC by individual cases. Synapses on spines have been sub-divided into those that are established on spine heads and those that are established on spine necks. We also differentiated between aspiny and spiny dendritic shafts. Data are given as percentages with the absolute number of synapses studied in parentheses. AS: asymmetric synapses; EC: entorhinal cortex; SS: symmetric synapses.

| EC layer | Case | Type of synapse | Synapses on spine heads | Synapses on spine necks | Synapses on aspiny dendritic shaft | Synapses on spiny dendritic shaft | Total synapses |
| --- | --- | --- | --- | --- | --- | --- | --- |
| II | AB1 | AS | 43.9% (118) | 0.7% (2) | 33.8% (91) | 21.6% (58) | 100% (269) |
|  |  | SS | 4.8% (1) | 4.8% (1) | 42.9% (9) | 47.5% (10) | 100% (21) |
|  | AB3 | AS | 39.3% (120) | 0.7% (2) | 29.8% (91) | 30.2% (92) | 100% (305) |
|  |  | SS | 13.9% (5) | 0% (0) | 27.8% (10) | 58.3% (21) | 100% (36) |
|  | IF10 | AS | 47.5% (122) | 0.4% (1) | 22.1% (57) | 30.0% (77) | 100% (257) |
|  |  | SS | 10.0% (2) | 0% (0) | 20.0% (4) | 70.0% (14) | 100% (20) |
|  | M16 | AS | 63.5% (207) | 0.9% (3) | 13.8% (45) | 21.8% (71) | 100% (326) |
|  |  | SS | 0% (0) | 0% (0) | 30.4% (7) | 69.6% (16) | 100% (23) |
| III | AB1 | AS | 53.8% (170) | 1.3% (4) | 15.2% (48) | 29.7% (94) | 100% (316) |
|  |  | SS | 22.6% (7) | 3.2% (1) | 16.1% (5) | 58.1% (18) | 100% (31) |
|  | AB3 | AS | 32.4% (110) | 0.3% (1) | 31.6% (107) | 35.7% (121) | 100% (339) |
|  |  | SS | 18.8% (6) | 6.2% (2) | 32.2% (10) | 43.8% (14) | 100% (32) |
|  | IF10 | AS | 48.7% (182) | 1.1% (4) | 20.0% (75) | 30.2% (113) | 100% (374) |
|  |  | SS | 0% (0) | 0% (0) | 23.1% (6) | 76.9% (20) | 100% (26) |
|  | M16 | AS | 52.4% (177) | 0.9% (3) | 15.1% (51) | 31.6% (107) | 100% (338) |
|  |  | SS | 9.7% (3) | 0% (0) | 19.3% (6) | 71.0% (22) | 100% (31) |

**Supplementary figure 1 (for Table 2).**
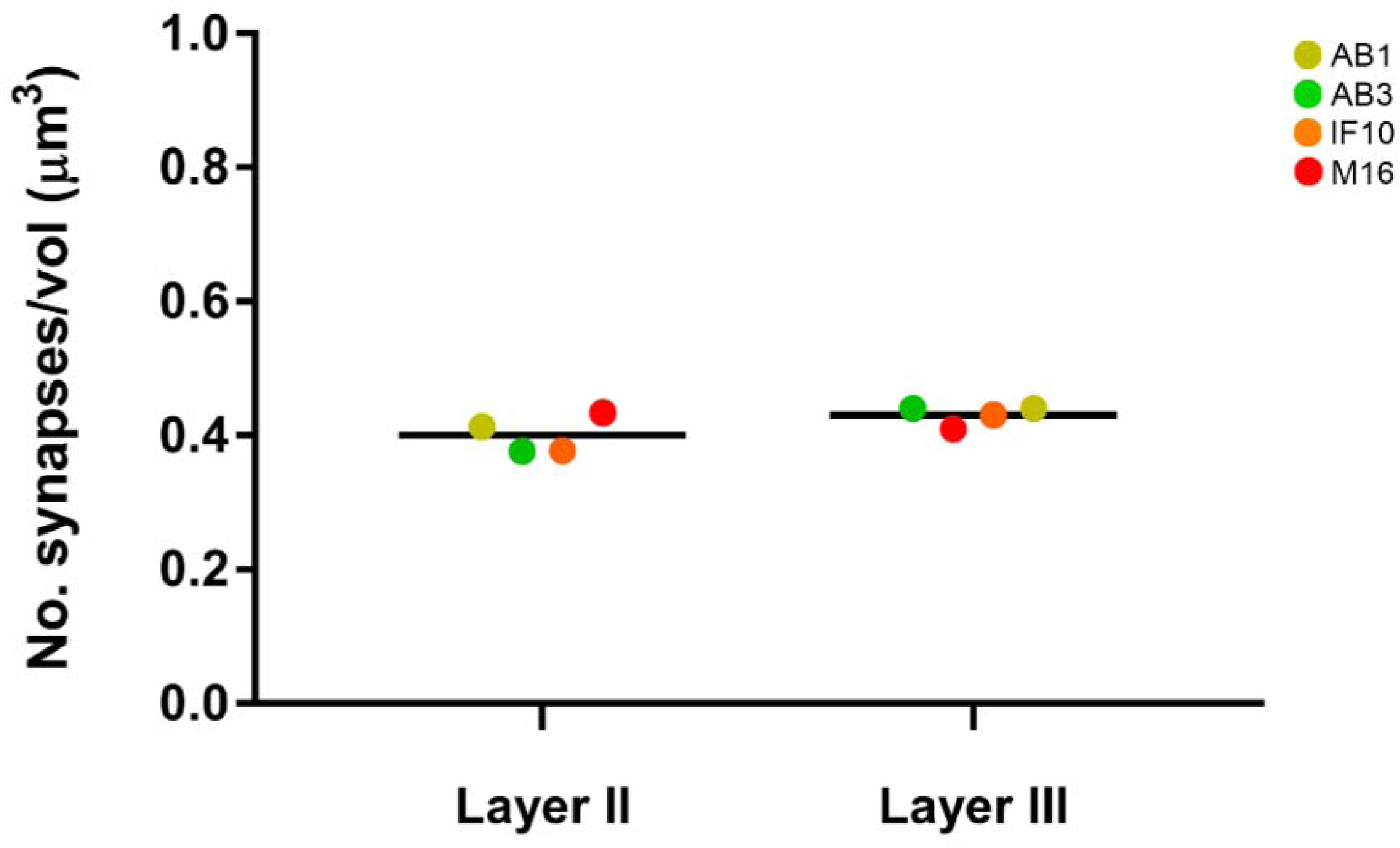
Graph showing the overall mean synaptic density in layers II and III of the EC. Each color corresponds to each case analyzed in the study, as denoted in the upper right-hand corner. No significant differences were found between layers (MW, p = 0.18).

**Supplementary figure 2.**
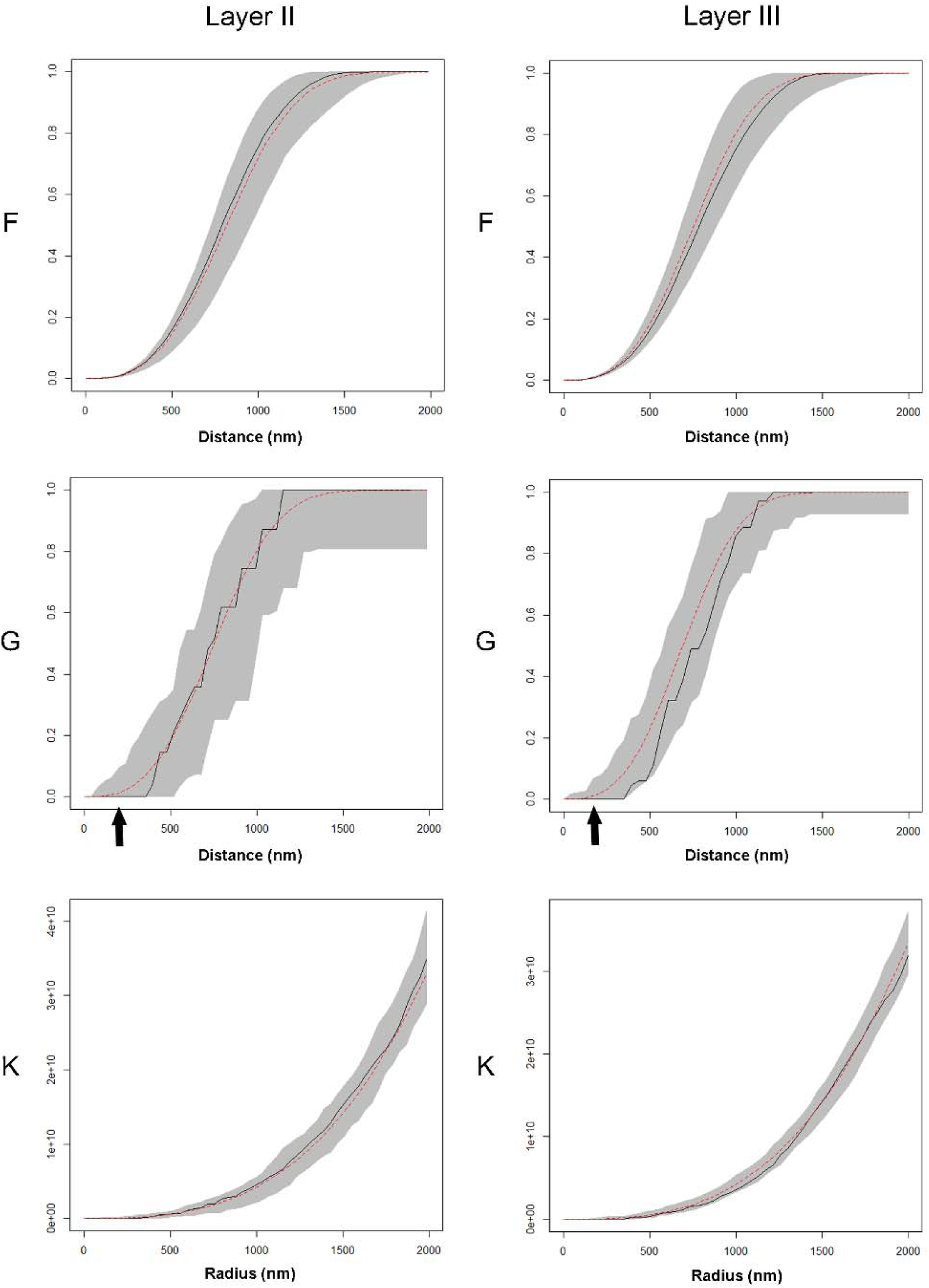
Analysis of the 3D synaptic spatial distribution in layers II and III of the EC. Red dashed traces correspond to a theoretical homogeneous Poisson process for each function. The black continuous traces correspond to the experimentally observed function. The shaded areas represent the envelopes of values calculated from a set of 99 simulations. All plots show a distribution which fits a Poisson function. In the G function, the arrow shows a dead space since there is a limit to hoe close synapses can be to each other since they cannot overlap in space. F: F function; G: G function; K: K function.

**Supplementary figure 3 (for Figure 5).**
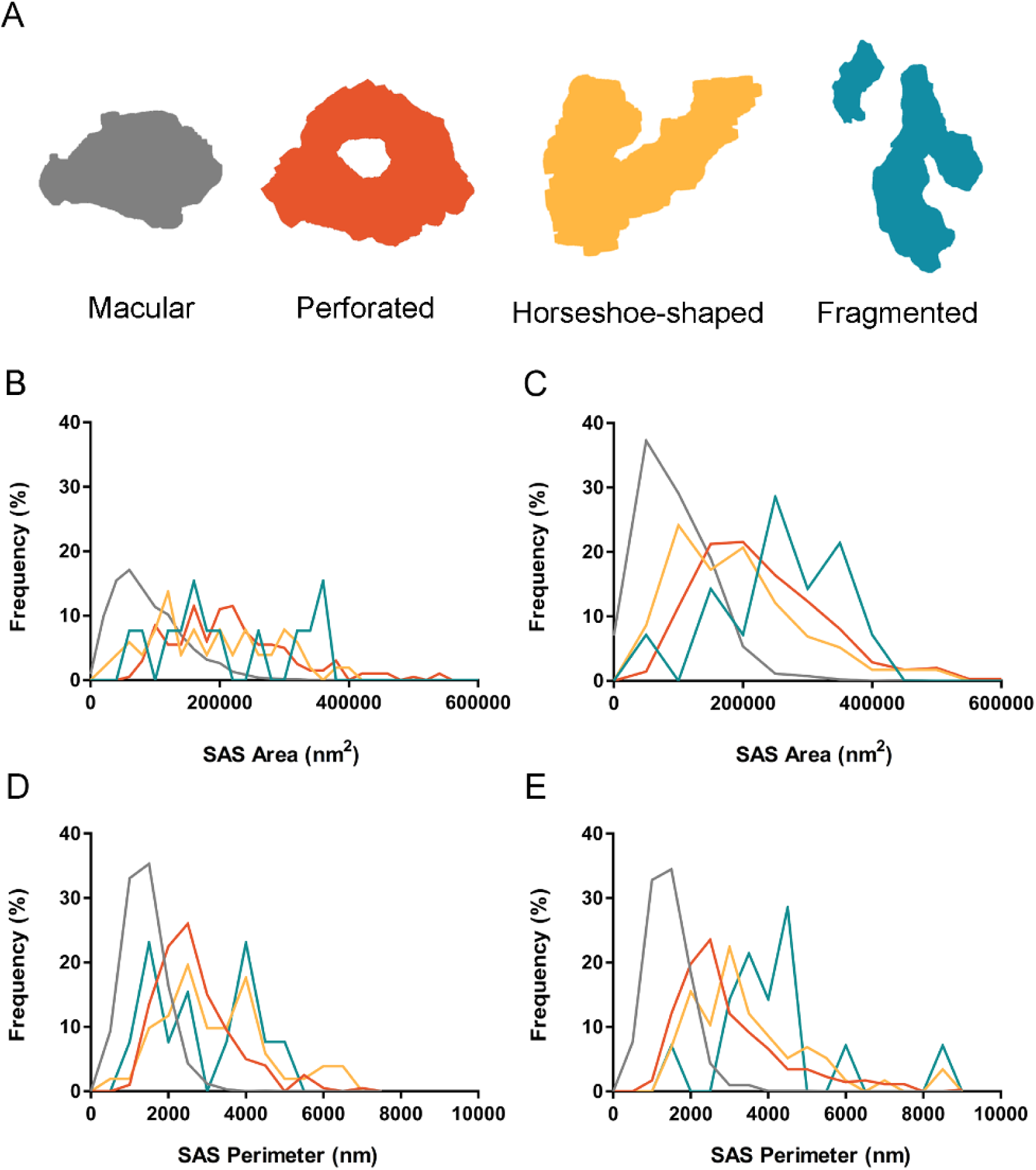
Schematic representation of the shape of the synaptic junctions: macular synapses, with a continuous disk-shaped PSD; perforated synapses, with holes in the PSD; horseshoe-shaped, with a tortuous horseshoe-shaped perimeter with an indentation; and fragmented synapses, with two PSDs with no connections between them **(A)**. Frequency histograms of the SAS area of macular, perforated, horseshoe-shaped and fragmented AS from layers II (**B)** and III **(C)** of the EC. Frequency histograms of the SAS perimeter of macular, perforated, horseshoe-shaped and fragmented AS from layers II (**D)** and III **(E)** of the EC. The mean SAS area and perimeter of macular synapses were significantly smaller than in perforated, horseshoe-shaped and fragmented synapses (ANOVA, p <0.001) in layers II and III of the EC.

**Supplementary figure 4 (for Figure 6).**
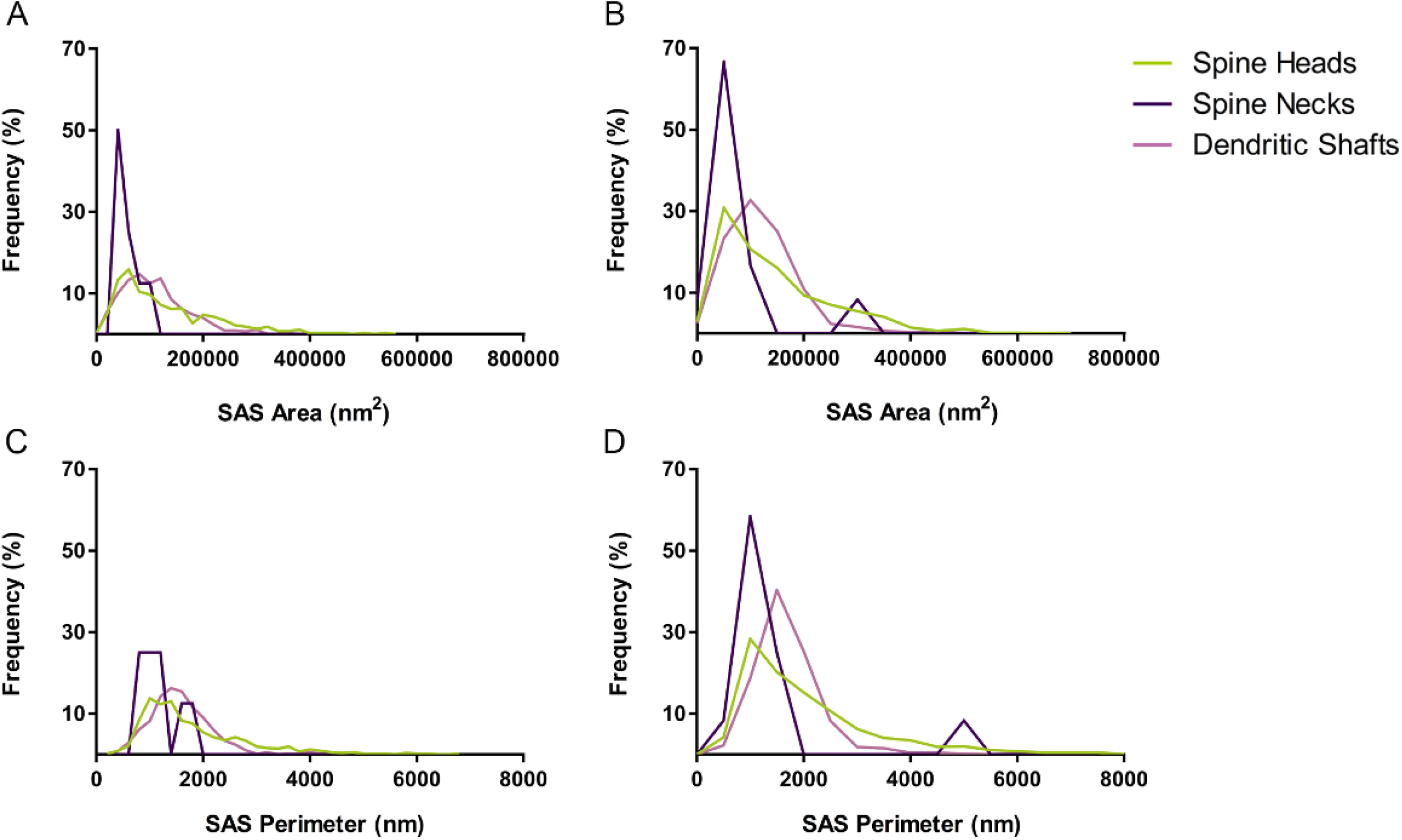
Frequency histograms of the SAS area **(A, B)** and perimeter **(C, D)** of AS targeting spine heads, spine necks and dendritic shafts in layers II **(A, C)** and III **(B, D)** of the EC. In the layer II samples, AS established on dendritic spine necks were smaller than AS established on dendritic heads or shafts (ANOVA, p <0.001).

